# Synthetic transcriptional control in the malaria parasite *Plasmodium falciparum*

**DOI:** 10.64898/2026.08.21.744319

**Authors:** Pablo Cárdenas Ramírez, Sebastian Smick, Sumanta Dey, Jacquin C. Niles

**Affiliations:** Department of Biological Engineering, Massachusetts Institute of Technology, Cambridge, Massachusetts, USA; R.F. Smith School of Chemical and Biomolecular Engineering, Cornell University, Ithaca, New York, USA; Pfizer, Inc., Cambridge, Massachusetts, USA; Broad Institute of MIT and Harvard, Cambridge, Massachusetts, USA

## Abstract

Malaria is responsible for over half a million deaths each year. However, our understanding of malaria parasite biology is hampered by a lack of molecular tools, particularly at the level of transcriptional control. In light of this, we have created two orthogonal systems for inducible transcriptional repression in the malaria parasite *Plasmodium falciparum* using bacterial repressor proteins. We achieve 200- to 800-fold repression of expression, improving on previous attempts at transcriptional regulation by two orders of magnitude and outperforming gold standard translational/post-transcriptional regulation systems. We developed automated DNA design software to apply this tool to conditional regulation of native gene expression, validating essentiality and chemogenetic interactions with both two parasite lipid kinases and *Pf*Kelch13, which is associated with artemisinin resistance. These tools can advance our understanding and engineering of malaria functional genomics, drug mechanisms, and gene regulation.

## TEXT

Malaria continues to be one of humankind’s deadliest infectious diseases. The most severe form of this mosquito-borne disease is caused by the eukaryotic parasite *Plasmodium falciparum* and results in around 600,000 deaths every year in some of the world’s most vulnerable communities^1^. Research on the parasite’s complex biology is hampered by the difficulty of implementing molecular tools with which to dissect gene function. Traditional functional genomics tools such as RNA interference (RNAi) and high-throughput CRISPR Cas9 knockout screens are not effective due to the low transfection efficiency of *P. falciparum*^2^ and absence of native RNAi^3^ and non-homologous end joining (NHEJ) DNA repair^4,5^ machinery.

To study gene function in *Plasmodium*, researchers modulate protein abundance and/or localization-specific function through inducible degradation^6,7^ or mislocalization^8^, translational/post-transcriptional regulation through recombinant RNAi^9^, inducible aptamers^10–13^ or ribozymes^14^, gene disruption through transposon mutagenesis^15^ or inducible DiCre-mediated knockout^16,17^, and transcriptional modulation through CRISPR interference (CRISPRi) using dCas9^18,19^ alone or fused to chromatin regulators^20,21^. For transcription specifically, the highest dynamic range achieved using CRISPRi is 1.5- to 10-fold, with variability across target genes^21^. A TetR-based transcriptional activation system systems for malaria parasites was first reported^22^ and then improved upon^23^ to achieve up to 16-fold regulation of genes expressed from the parasite genome. Although the system has seen use, particularly in the rodent malaria model *Plasmodium berghei*^24–29^, translational control tools remain the most widely adopted methods for modulation of gene expression in malaria parasites due to their higher reliability across genetic contexts and increased dynamic range. The highest dynamic range among these is achieved by the TetR-DOZI-aptamer system, which enables inducible, reversible, and tunable control of expression with a dynamic range exceeding 100-fold^11,13^ and has seen use in dozens of studies of *P. falciparum*, as reviewed previously^30^. Thus, no tool is currently capable of reliable, reversibly tunable, and high-dynamic range transcription al control.

The lack of efficient tools for transcriptional control in *Plasmodium* is significant given that transcription al regulation is critical in maintaining tightly-controlled gene expression across the parasite’s life cycle. In addition to well-documented translational/post-transcriptional^31^ and epigenetic mechanisms^32^, mRNA expression is tightly regulated throughout the 48 h intraerythrocytic development cycle (IDC)^33–40^ during which chromatin remains mainly in an open state^37–40^. Furthermore, important aspects of parasite biology such as variable surface antigen expression appear to depend on noncoding RNA expression^32,41^. These findings highlight the importance of transcriptional regulation for parasite biology and the potential impact of tools that reliably control it.

In this study, we engineer two synthetic systems achieving transcriptional control in *P. falciparum*. Using nuclear-localized bacterial repressor proteins and arrays of their corresponding DNA operator sequences, we achieve titratable, modular repression across promoter contexts with different profiles of expression strength timing across the IDC timing ad strength. The most effective of these systems achieves 200- to 800-fold dynamic regulatory range and is active across the entire IDC. We apply this system to confirm the essentiality of two lipid kinases (*Pf*PI3K and *Pf*PI4K) and a modulator of artemisinin resistance, *Pf*Kelch13. We also establish utility of these systems in translational drug discovery-related studies through validating several compound-gene interactions.

## Results

### Engineering transcriptional control in malaria parasites through DNA operator arrays

We hypothesized that by placing enough DNA operator sequences between a *P. falciparum* 5’ untranslated region (UTR; expected to contain the transcriptional promoter element) and its coding sequence, we could create enough steric hindrance for the RNA polymerase II (pol II) transcription complex to decrease transcription of a target gene. This has been done in mammalian cells^42–46^ as well as in other eukaryote parasites^47^ with the bacterial repressor TetR, which binds the *tetO* operator sequence in the absence of tetracycline and its less toxic analogue anhydrotetracycline (aTc). We repurposed plasmid pSG372^13^ to constitutively express TetR and assess transcriptional regulation of a firefly luciferase (FLuc) reporter gene based on this principle (Fig. 1A). Increasing numbers of *tetO* sequences were amplified from an existing plasmid^48^ and inserted between the FLuc coding sequence and the well-characterized pCAM 5’ UTR sequence from the *P. falciparum* calmodulin (PF3D7_1434200) 5’ UTR^49^. Constitutive expression of an orthogonal reporter *Renilla* luciferase (RLuc) provided a normalization reference for detecting changes in FLuc expression.

**Figure 1.**
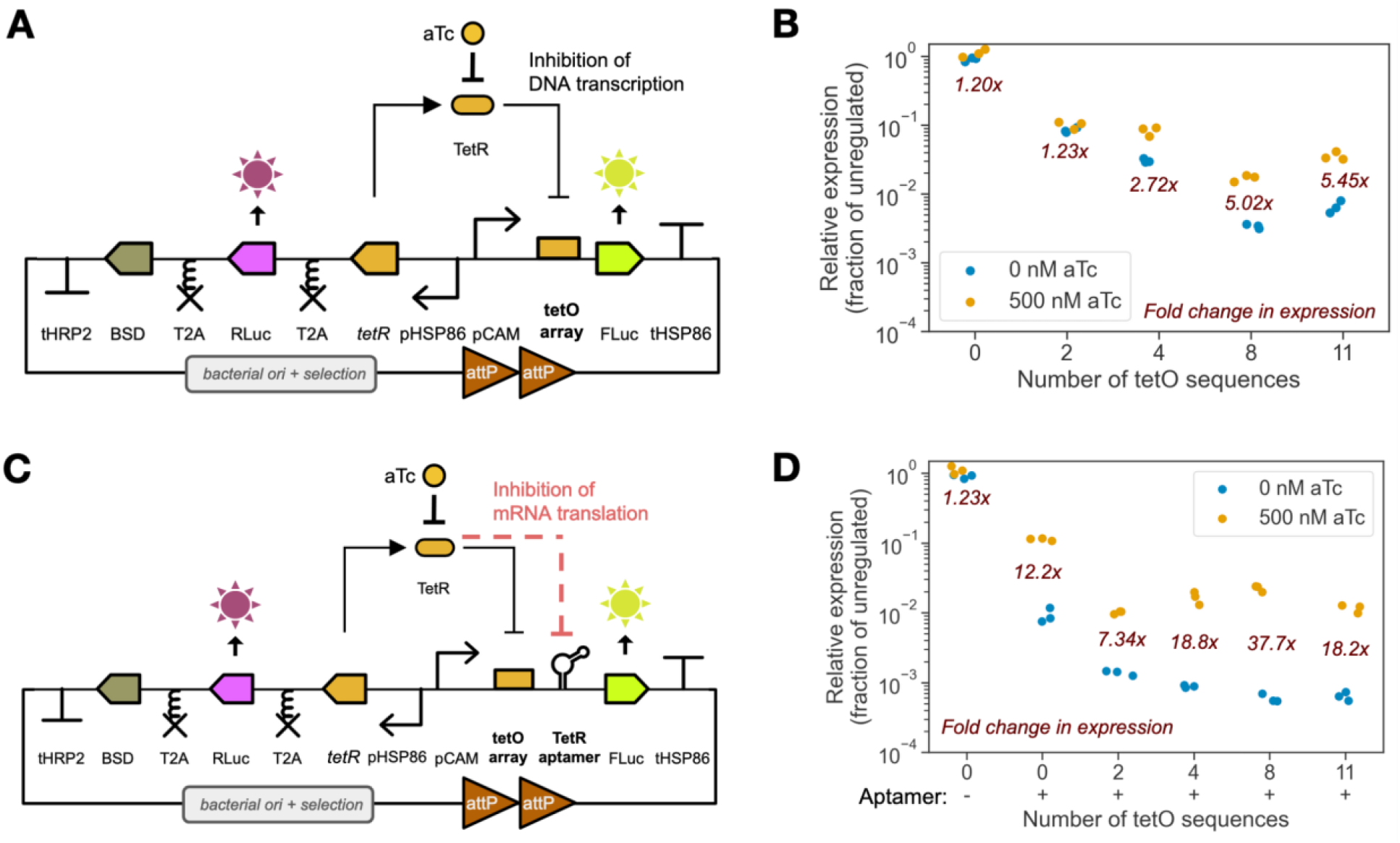
Bacterial repressor-operator interactions can modulate gene transcription in *Plasmodium falciparum*. **(A)** A circular plasmid used for *attP-attB* integration into parasite genomes serves as a testbed for regulation of expression of a target reporter gene, firefly luciferase (FLuc), using increasing numbers of *tetO* operators that bind to repressor TetR in the absence of anhydrotetracycline (aTc). TetR is expressed constitutively alongside the orthogonal reporter *Renilla* luciferase (RLuc) and a blasticidin-S deaminase (BSD) selection marker, separated by self-cleaving T2A peptides. Transcription of FLuc in *P. falciparum* is achieved using the 5’ and 3’ untranslated regions (UTRs) of calmodulin (pCAM) and HSP86 (tHSP86), respectively, while transcription of the *tetR*-RLuc-BSD unit was achieved with the 5’ and 3’ UTRs of HSP86 (pHSP86) and HRP2 (tHRP2), respectively. **(B)** Relative expression of FLuc and fold change in expression under induction by aTc for increasing numbers of *tetO* repeats in the operator array. **(C)** An RNA aptamer sequence capable of aTc-dependent binding to TetR combines translational repression to the previous transcriptional regulation. **(D)** Relative expression of FLuc and fold change in expression under induction by aTc for increasing numbers of *tetO* repeats in the operator array with a downstream TetR aptamer sequence.

Using this setup, we confirmed that using two operators did not achieve significant changes in FLuc expression. However, with ≥2 *tetO* operators, we observed aTc-dependent regulation of FLuc expression, which peaked at ≈5-fold (Fig. 1B). Increasing the number of *tetO* repeats also decreased maximum FLuc expression relative to the no operator reference. We also tested the effect of combining TetR-dependent transcriptional and TetR aptamer-based post-transcriptional regulation^10,13^. This reduced background (-aTc) FLuc expression by ≈10-fold relative to the context with only transcription repression, and achieved a dynamic range > 18-fold for >2 *tetO* repeats (Fig. 1D).

### A TetR-based, aTc-inducible transcriptional regulator with 20-fold dynamic range

We next sought to improve the regulatory dynamic range of the TetR-*tetO* system through different approaches. We reduced the length of operator array spacers to 2–3 bp between each repeat to reduce separation between the 5’ UTR and start codon, then added an N-terminal SV40 nuclear localization signal (NLS) onto TetR to increase its concentration in the nucleus. While this increased fold regulation (Fig. S1A), it also increased toxicity to the parasites (Fig. S1B–D). In fact, parasites transfected with SV40–TetR were recovered only when cultures were maintained with aTc present, implicating SV40–TetR binding to chromosomally-integrated *tetO* repeats as the likely culprit. Supporting this interpretation, aTc removal from these cultures inhibited parasite replication (Fig S1G); additionally, parasites harboring SV40–TetR but no genomic *tetO* repeats or vice versa could be recovered in the absence of aTc and exhibited no aTc-dependent replication defects (Fig S1G). We resolved the toxicity issue by substituting the strong HSP86 5’ UTR used originally to drive SV40– TetR expression with the weaker mRPL2 5’ UTR (pmRPL2; Fig. 2A)^50,51^. Reduced SV40–TetR expression preserved effective regulation of FLuc and restored normal parasite replication in the absence of aTc (Figs. S1B, S1C, S1E). By quantitative RT-PCR, we observed a 12-fold change in FLuc mRNA abundance upon aTc induction (Fig. 2B), confirming effective transcriptional regulation. Over a range of aT c concentrations, w e established that for both the No-NLS^*hsp86*^ and SV4 0–TetR^mRLP2^ regulated contexts, FLuc expression is titratable (Fig. 2C). Basal “leaky” expression is ≈6-fold lower in the latter context, and both exhibit similar fully induced F L uc expression levels. Thus, the dynamic range s for the No-NLS-TetR^*hsp86*^ and SV40–TetR^mRLP2^ systems are ≈5- and ≈20-fold, respectively, with the improved performance due primarily to a reduction in leaky expression with the nuclear-localized repressor. We also confirmed stable regulation is maintained over at least two replication cycles (Fig. 2D).

**Figure 2.**
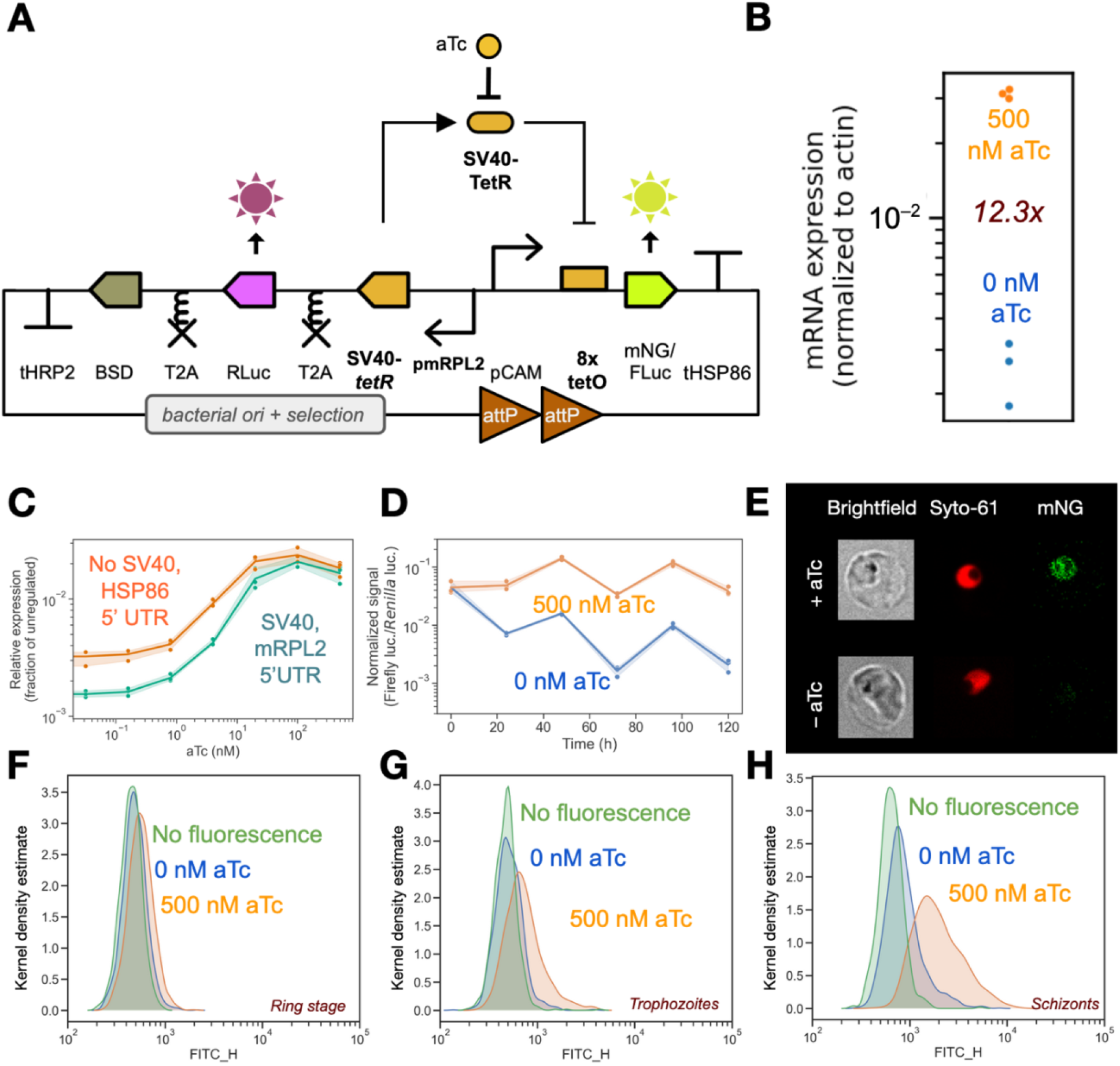
A TetR-based, aTc-inducible system for transcriptional control in *Plasmodium falciparum*. **(A)** TetR repressor fused with an SV40 nuclear localization signal (NLS) is expressed from a weak mRPL2 5’ untranslated region (UTR) containing the promoter (pmRPL2). SV40_TetR binds an array of 8 *tetO* operator repeats in an anhydrotetracycline-(aTc-) inducible manner, regulating expression of either mNeonGreen (mNG) or firefly luciferase (FLuc) from a calmodulin 5’ UTR and promoter (pCAM). Consitutively expressed *Renilla* luciferase (RLuc) is used to normalize FLuc expression, while blasticidin-S deaminase (BSD) serves as a selection marker. T2A ‘skip’ peptides allows for expression of multiple proteins encoded by a single transcript, and 3’ UTR regions containing terminators from the *hrp2* and *hsp86* genes (tHRP2, tHSP86) complete the transcriptional units. **(B)** RT-qPCR measurement of FLuc mRNA abundance (normalized to actin) comparing exposure to 0 nM and 500 nM aTc for 72 h. **(C)** Relative FLuc expression after 72 h of exposure to increasing concentrations of aTc when expressing TetR with a strong HSP86 5’ UTR and no nuclear localization or expressing SV40-fused TetR with a weak mRPL2 promoter. **(D)** Normalized FLuc expression signal measured across multiple 48 h parasite intraerythrocytic development cycles (IDCs). **(E)** Expression of mNG from pCAM in late-stage parasites observed by fluorescence microscopy after 72 h of growth in presence or absence of aTc. Expression of mNG after 72 h of growth in presence and absence of aTc, measured through flow cytometry in the fluorescein (FITC) channel, for **(F)** ring stage parasites, **(G)** trophozoites, and **(H)** schizonts.

To confirm promoter context-independent regulation is attainable, we used RNA Seq expression data^52,53^ to select several 5’ UTRs with different temporal and peak expression profiles during the replication cycle to test in our reporter system. These included the chloroquine resistance transporter (CRT, PF3D7_0709000; 1463 bp), a weak 5’ UTR most active during ring stages; elongation factor 1-alpha (EF1α, PF3D7_1357000, 806 bp), a medium strength 5’ UTR most active in late rings and early trophozoites; calmodulin (CAM, PF3D7_1434200, 624 bp), a medium strength 5’ UTR most active in late trophozoites and schizonts; heat shock protein 70 (BIP, PF3D7_0917900, 1949 bp), a strong 5’ UTR most active in trophozoites and schizonts; and apical membrane antigen 1 (AMA1, PF3D7_1133400, 1448 bp), a strong 5’ UTR with activity tightly restricted to schizonts and merozoites. Using an array of 8 *tetO* repeats, we observed robust regulation of FLuc expression (15– 52-fold) for all 5’ UTRs active in trophozoite and schizont stages. We found reduced regulation (1.8-fold) with the ring stage 5’ UTR, CRT (Fig. S2), possibly explained by low TetR expression early in the life cycle due to the expression pattern of the mRPL2 5’ UTR^52,53^. We also confirmed that regulation is target gene independent by substituting FLuc for the mNeonGreen (mNG) reporter and measuring fluorescent signal by flow cytometry and fluorescence microscopy. In the 5’-C AM UTR context, we observed fluorescence signal during live cell fluorescence microscopy under induction by aTc, and background levels in its absence (Fig. 2E). We confirmed these results using flow cytometry (Figs. 2F–H, S3C, S3E). It is worth noting that fluorescence intensity in ring stage parasites is challenging to distinguish from background fluorescence due to their small size and background autofluorescence. Nevertheless, flow cytometry confirmed that control of expression is tunable by aTc at the single-cell level in late-stage parasites (Figs. S4A, S4B), as we had observed in bulk parasite culture using a luminescence reporter (Fig. 2C).

### A LacI-based, IPTG-inducible transcriptional regulator with 400-fold dynamic range

To examine whether other inducible bacterial repressors can regulate transcription in *P. falciparum*, we used the same plasmid system used for TetR (Fig. 1A) and replaced the TetR coding and *tetO* array sequences with three other repressor-operator pairs inducible by different small molecules, chosen for their use in other synthetic biology contexts. Two failed to function effectively due to toxicity of the inducer (DAPG, cumate) or repressor expression (no CymR repressor transfectants were recovered after three attempts) (Figs. S5A, S6). However, the LacI inducer isopropyl β-D-1-thiogalactopyranoside (IPTG) was well-tolerated, even up 5 mM) (Fig. S6), and our initial LacI-*lacO* system design achieved a dynamic range of 14-fold, Fig. S5A). This was surprising given that amplicon sequencing of transfected parasites revealed that 5 out of 8 *lacO* operator sequences had been lost, presumably due to recombination. Although we encountered no DNA stability issues with *tetO* arrays, direct *lacO* repeats proved unstable, requiring assembly and growth in *Escherichia coli* at 28 ºC instead of 37 ºC prior to transfection. To improve the stability of the *lacO* DNA array, we reverted to using spacers of different lengths (7–27 bp) and variations of the two wild-type *lacO* sequences found in nature as well as the optimized, synthetic *Osys* operator sequence^54–56^. We used these variants to edit the sequences found in a previous synthetic *lacO* array that had been optimized for DNA stability^57^ to create mixed arrays of 4 and 8 *lacO* variant repeats, replacing our original, unstable array of direct repeats. The improved arrays were stable during bacterial growth at 37 ºC and remained intact following transfection into *P. falciparum* (Fig. S7), achieving 182- and 409-fold regulation of FLuc expression, respectively (Fig. S5A). Furthermore, SV40_LacI expressed from the strong *hsp86* 5’ UTR and its inducer IPTG appear to be non-toxic (Figs. S5B–F).

We selected the 8-repeat *lacO* array with its higher dynamic range (Fig. S5A) for further assessment within the genetic context illustrated in Fig. 3A. The magnitude of regulation was large enough to result in considerable changes in mRNA expression after 72 h (Fig. 3B). As with the TetR-based system, regulation was found to be tunable across the full dynamic range according to IPTG concentration. However, the LacI-based system permitted a higher expression level in its induced state, reaching 43% of the maximum expression of unregulated FLuc with no operators (Fig. 3C). In contrast, the SV40_TetR system achieved 2.1% of maximum unregulated expression, while the 5’ TetR aptamer translational control system achieved a figure of 13%^10,13^. Despite the fact that the induced expression state is higher, LacI-based control also achieved a lower expression level when not induced, resulting in less leaky expression: 0.098% of maximum unregulated expression (Fig. 3C), compared to 0.12% and 1.0% for the SV40_TetR and 3’ TetR aptamer system^10,13^, respectively. Regulation was once again preserved across IDCs (Fig. 3D). Notably, strong repression was observed for all five 5’ UTR regions with promoters tested (212- to 830-fold regulation), including that of CRT (Fig. S8). Control of expression was easily visible by fluorescence microscopy when the FLuc coding sequence was replaced by that of mNG (Fig. 3E). Robust control of expression in all RBC life stages, including rings, was verified using flow cytometry (Figs. 3F–H, S3D, S3F). Continuous, tunable variation in expression at the single-cell level was also observed (Figs. S4C, S4D).

**Figure 3.**
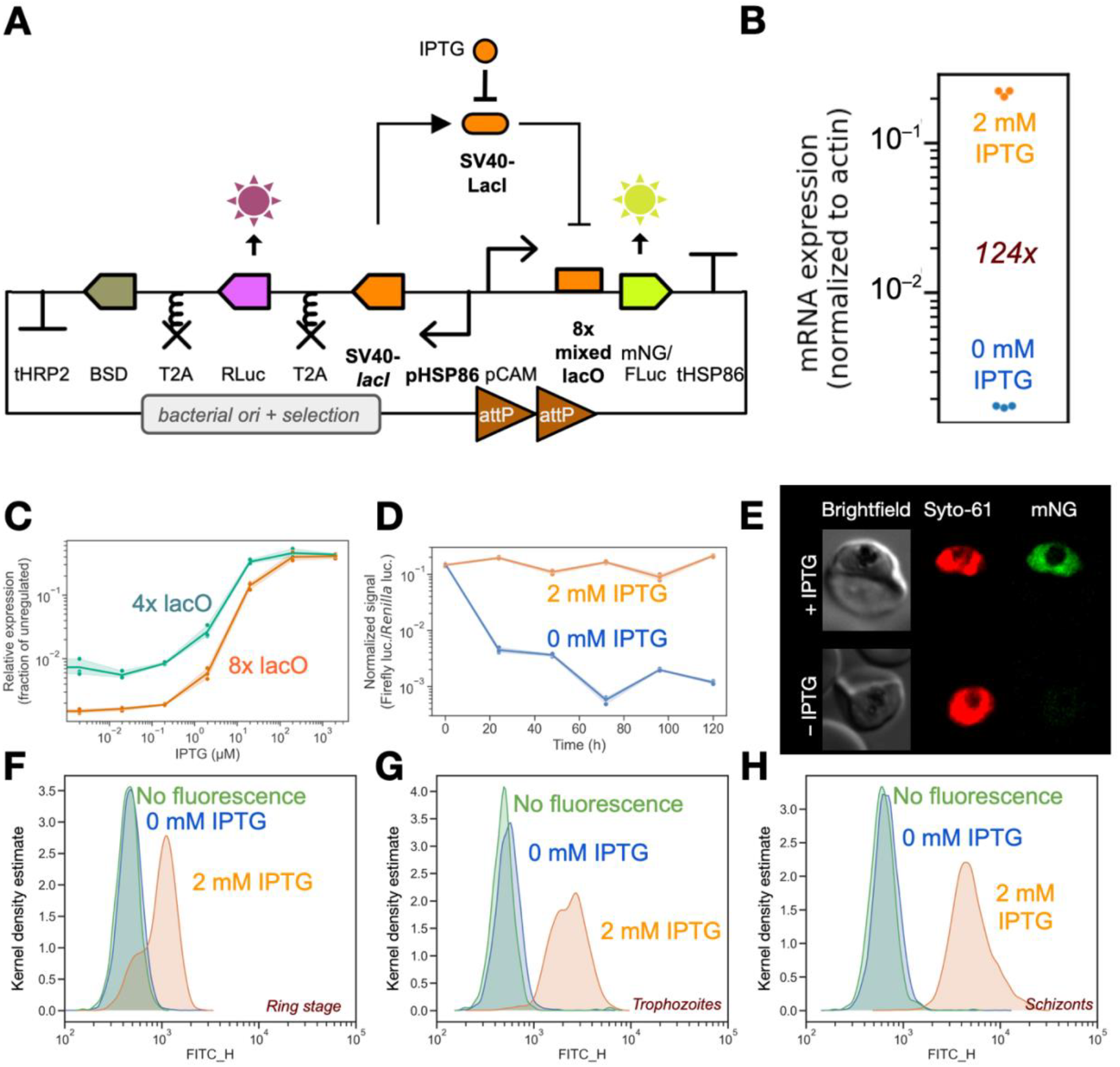
A LacI-based, IPTG-inducible system for transcriptional control in *Plasmodium falciparum*. **(A)** LacI repressor tagged with nuclear localization signal (NLS) SV40 is expressed from a strong HSP86 5’ untranslated region (UTR) and promoter (pHSP86). SV40_LacI then binds to an array of 8 *lacO* operator variant repeats in an isopropyl β-D-1-thiogalactopyranoside-(IPTG-) inducible manner, regulating the expression of either mNeonGreen (mNG) or firefly luciferase (FLuc) from a calmodulin 5’ UTR and promoter (pCAM). *Renilla* luciferase (RLuc) allows for a measurement of constitutive expression to use in normalizing FLuc expression, while blasticidin-S deaminase (BSD) serves as a selection marker. Self-cleaving T2A peptides separate independent proteins on the same coding sequence, and 3’ UTR regions containing the terminators from genes HRP2 and HSP86 (tHRP2, tHSP86) complete the transcriptional units. Regulation of expression is observed both with **(B)** FLuc mRNA abundance by RT-qPCR (normalized to actin) after 72 h of growth in the presence or absence of IPTG. **(C)** FLuc expression under increasing concentrations of IPTG after 72 h of growth using constructs containing either 4 or 8 *lacO* variant repeats. **(D)** Normalized FLuc expression signal measured across multiple 48 h parasite intraerythrocytic development cycles (IDCs). **(E)** Expression of mNG from pCAM observed by fluorescence microscopy after 72 h of growth with and without IPTG. Expression of mNG at 72 h of growth in presence and absence of IPTG, measured by flow cytometry in the fluorescein (FITC) channel, for **(F)** ring stage parasites, **(G)** trophozoites, and **(H)** schizonts.

In view of the success of LacI-based regulation in *P. falciparum* using *lacO* operators at the start of the coding sequence, we attempted to modulate FLuc expression using an array of 8 *lacO* variant operators at the 3’ end of the coding sequence, inserted between the stop codon and 3’ UTR rather than upstream of the 5’ ATG. 3’ UTRs are important elements for correct mRNA processing, and we hypothesized that interrupting transcription at this point could still afford strong inhibition of expression, even if the coding sequence had already been transcribed. Although this 3’ targeting strategy still achieved substantial regulation of expression (7.4-fold, Fig. S5A) with a very high level of induced expression, the repressed state had much higher leakiness and the magnitude of regulation was thus nearly 50-fold lower than its 5’ targeting counterpart.

### Robust transcriptional knockdown validates lipid kinase essentiality and compound interactions

With this new tool for conditional expression in *P. falciparum*, we next attempted to regulate the expression of native genes. We selected the lipid kinases phosphatidylinositol 3-kinase (*Pf*PI3K, PF3D7_0515300) and phosphatidylinositol 4-kinase beta (*Pf*PI4K, PF3D7_0509800), implicated in lipid signaling during processes such as endocytosis and cytokinesis^58,59^. Both *Pf*PI3K and *Pf*PI4K are expected to be essential genes based on the results of transposon mutagenesis screening^15^ and directed attempts to knock out each gene^59,60^ all failing to disrupt either coding sequence. However, earlier attempts at using conditional knockdown techniques such as the TetR_DOZI-aptamer system^10,13^ failed to demonstrate each gene’s essentiality, with parasites maintaining growth in the knocked down state, which we confirmed with our own experiments (Fig. S9). We hypothesized this was due to leaky expression of each gene and could therefore be remedied by using a system with more stringent control of expression.

To this end, we designed a linear plasmid system housing the LacI-*lacO* system and allowing its integration into any target gene through Cas9-mediated homologous recombination. This plasmid, pPC101 (Fig. 4A), follows the logic of previous pJAZZ-based linear vector systems^13^, and depends on either cotransfection with a pCRISPR plasmid already expressing Cas9 and T7 polymerase or single transfection into a parasite line already expressing both proteins. The selection process for the different DNA components required for gene editing (single guide RNA, left and right homology regions, and recoded region), along with the design of all relevant oligo and gene fragment sequences for DNA assembly, is now automated by selecting the new “pPC101” option in GeneTargeter, an online *Plasmodium* DNA designer tool developed previously^30^ (www.genetargeter.mit.edu).

**Figure 4.**
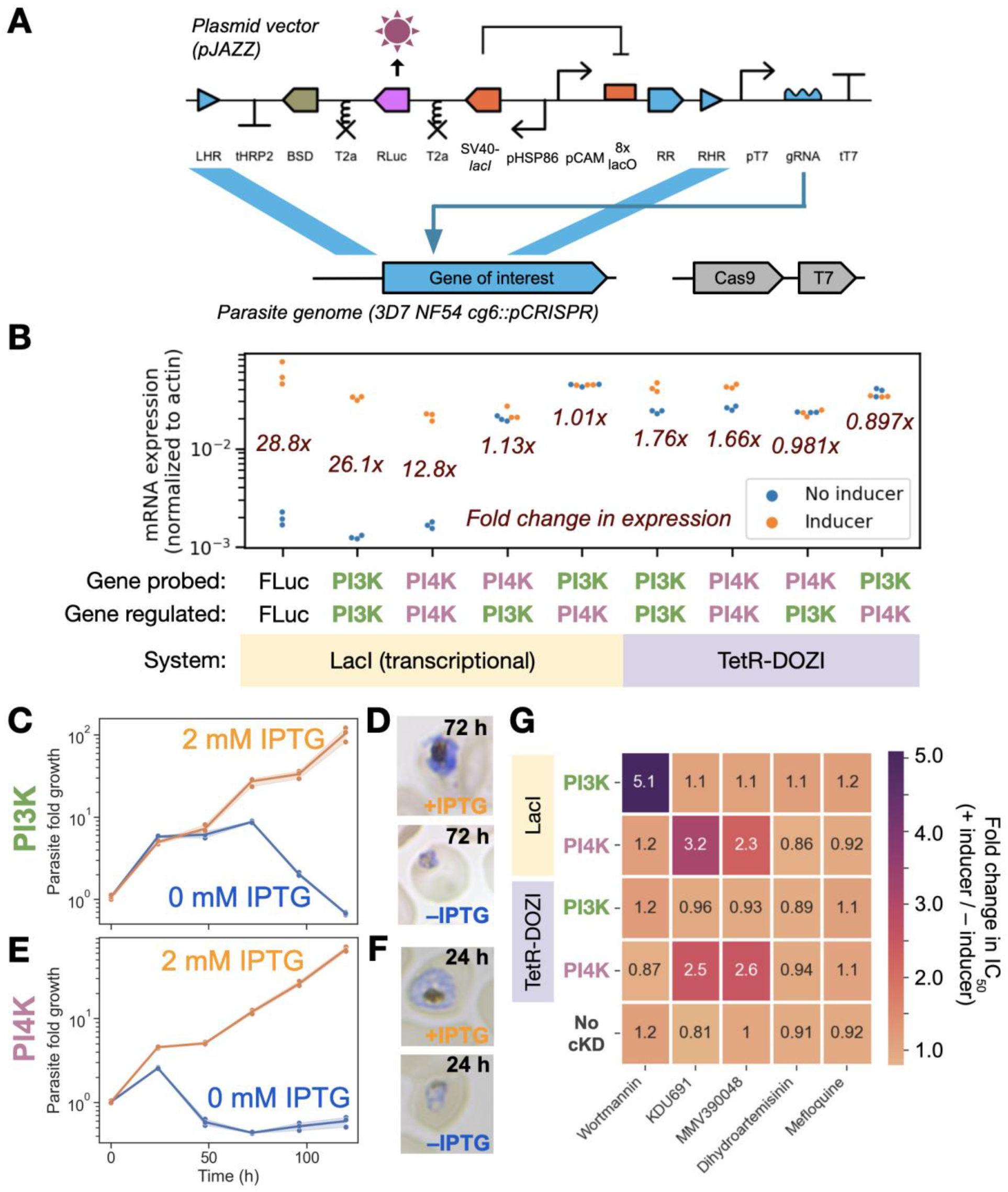
Conditional regulation of transcription validates the essentiality of *P. falciparum* phosphatidylinositol 3-kinase (*Pf*PI3K) and phosphatidylinositol 4-kinase beta (*Pf*PI4K). **(A)** The pPC101 linear plasmid system for conditional regulation of native gene transcription in *P. falciparum* uses Cas9-mediated homologous recombination (HR) to insert a LacI-*lacO* regulation system into a target coding sequence. pPC101 expresses a single guide RNA (sgRNA) using a T7 polymerase promoter (pT7) and terminator (tT7), which leads Cas9 to cause a double-stranded break at the gene of interest. The break is later repaired with left and right homology regions (LHR and RHR, respectively) on the plasmid, resulting in the regulatory system being integrated at the 5’ end of the gene and part of the gene’s sequence being reconstituted with a recoded region (RR). **(B)** RT-qPCR (normalized to actin) 24 h after IPTG removal shows mRNA expression for targeted and off-target genes in late-stage parasites. **(C)** Parasite growth, measured as fold change in RLuc luminescence, following knockdown of *Pf*PI3K by removal of isopropyl β-D-1-thiogalactopyranoside (IPTG). **(D)** Parasite morphology after 72 h of growth with or without IPTG for *Pf*PI3K knockdown. **(E)** Parasite growth following knockdown of *Pf*PI4K due to IPTG removal. **(E)** Parasite morphology after 72 h of growth with or without IPTG for *Pf*PI4K knockdown. **(G)** Fold change in IC_50_ of antimalarial compounds, including putative and known inhibitors of *Pf*PI3K (wortmannin) and *Pf*PI4K (KDU691, MMV390048), when using LacI-based transcriptional knockdown or TetR-DOZI translational knockdown of *Pf*PI3K or *Pf*PI3K..

We used designs generated with the help of GeneTargeter to construct pPC101-type vectors targeting *P. falciparum Pf*PI3K and *Pf*PI4K for conditional regulation of transcription. These constructs were transfected in the presence of IPTG into parasites with genome-integrated Cas9 and T7 expression. Transfected parasites were recovered easily in both cases. Although both proteins were tagged with HA epitopes, western blots using anti-HA antibodies did not reveal the presence of wither, possibly due to their large size and membrane involvement. Nevertheless, the transcriptional nature of our knockdown technique allows us to quantify transcript levels as an alternative validation method. Therefore, levels of *Pf*PI3K and *Pf*PI4K transcripts was assessed by reverse transcriptase quantitative polymerase chain reaction (RT-qPCR) 24 h after knockdown while parasite populations were still not fully depleted. Knockdown was found to be gene-specific, comparable to that observed with FLuc, and able to achieve levels of transcript abundance similar to wild-type expression when induced by IPTG (Fig. 4B). The greatest fold change in expression was seen at different times for each gene (Fig. S10), reflecting differences in mRNA degradation kinetics across the life cycle^61,62^. TetR-DOZI regulation of *Pf*PI3K and *Pf*PI4K resulted in measurable changes in transcript abundance, as noted by others^63,64^. It is worth noting that native transcript abundance was more closely achieved by the induced LacI system than the induced TetR-DOZI system, the latter of which resulted in transcript levels higher than those of native expression (Fig. 4B). Luminescence-based growth assays and microscopy confirmed both *Pf*PI3K and *Pf*PI4K as essential. *Pf*PI3K knockdown had no visible effect in the first IDC after IPTG removal, but parasite growth stagnated by 72 h (Fig. 4C) with parasites failing to develop into the late stages (Fig. 4D) and was then followed by cell death. Knockdown of *Pf*PI4K had much faster effects, resulting in an immediate decrease in parasite growth as early as 24 h (Figs. 4E, 4F) and rapid cell death thereafter.

Having validated these conditional knockdown lines for *Pf*PI3K and *Pf*PI4K, we next used them to determine compound-target interactions for these two genes. Conditional knockdown dose response assays are useful tools to establish whether a compound inhibits a specific protein, thanks to the fact that decreasing the expression of a given protein will reduce its available supply within the cell and hypersensitize the cell to compounds that target the inhibited protein. This is visualized as a leftwards shift in the half maximal inhibitory concentration (IC_50_) of the compound in a dose response curve when the target gene is knocked down^59^. We carried out these assays using wortmannin, a classical *Pf*PI3K inhibitor in mammalian cells^65^ that had not been seen to inhibit *Pf*PI3K in *P. falciparum*, as well as KDU691 and MMV390048, two compounds previously identified as *Pf*PI4K inhibitor through other methodologies^59,66^. Dihydroartemisinin (DHA) and mefloquine were included as control compounds targeting a different aspect of parasite physiology. By inhibiting the expression of each lipid kinase and quantifying growth across a range of concentrations for each compound, we confirmed wortmannin as a *Pf*PI3K inhibitor in malaria parasites (Figs. 4G, S11A) and recapitulated targeting of *Pf*PI4K by both KDU691 and MMV390048 (Figs. 4G, S11G–H). These interactions were found to be specific, as KDU691 and MMV390048 did not target *Pf*PI3K (Figs. 4G, S11B–C) and wortmannin was not found to target *Pf*PI4K (Figs. 4G, S11F). DHA and mefloquine had no interaction with either kinase (Figs. 4G, S11D–E, S11I–J). TetR-DOZI conditional knockdown of *Pf*PI3K and *Pf*PI4K was able to capture the interactions between *Pf*PI4K and its inhibitors KDU691 (Figs. 4G, S11Q) and MMV390048 (Figs. 4G, S11R), but not the interaction between *Pf*PI3K and wortmannin (Figs. 4G, S11K). A parasite line with no knockdowns showed no IC_50_ shifts denoting interactions with any compound, as expected (Figs. 4G, S11U–Y).

It is worth noting that the *Pf*PI4K knockdown line was revealed to be hypomorphic, given that even fully induced expression of *Pf*PI4K resulted in a shift in IC_50_ for *Pf*PI4K inhibitors in the knockdown line (Figs S11G–H) when compared to lines with wild-type *Pf*PI4K expression (Figs S11B–C, L–M, V–W). This is likely due to the different expression patterns of the calmodulin 5’ UTR pCAM, used in pPC101 to drive expression of the gene being targeted for regulation, and the native *Pf*PI4K 5’ UTR it replaced. Conversely, TetR-DOZI knockdown led to an increase in *Pf*PI4K expression (Figs S11Q–R) over wild-type, possibly due to modified translational dynamics occurring with the replacement of the gene’s 3’ UTR that pSN054 TetR-DOZI modification implies^13^. Regardless, the LacI-based regulatory system was still able to show an IPTG-dependent IC_50_ shift for both *Pf*PI3K and *Pf*PI4K inhibitors.

### Reversible transcriptional knockdown reveals PfKelch13 compound interactions

Resistance to artemisinin in malaria parasites was first associated with mutations in the gene encoding Kelch propeller protein *Pf*Kelch13 through field isolates. Resistance in the field manifests as delayed parasite clearance when faced with temporary exposure to artemisinin^67^. However, when these clinically resistant parasite isolates are subjected to dose response curves and growth assays with constant drug exposure, they behave similarly to parasites with full sensitivity to^67–69^. Because of this, carefully-timed drug exposure assays such as the ring stage survival assay (RSA) have been developed to study resistance to artemisinin *in vitro*^67,70–72^. The crucial role of the timing of *Pf*Kelch13 expression and artemisinin exposure makes this system an attractive proving ground for our new tools for transcriptional regulation. Furthermore, multiple lines of evidence have been necessary to finally ascertain *Pf*Kelch13’s role in drug resistance, in which mutants reduce hemoglobin endocytosis and thus reduce pro-drug conversion by free heme for a limited amount of time^73^. We thus asked if temporary conditional knockdown could uncover the relationship between *Pf*Kelch13 and artemisinin resistance where constant drug exposure and gene expression techniques had failed.

We therefore applied the LacI-based transcriptional control system to the regulation of *Pf*Kelch13. Following the same design procedure outlined for *Pf*PI3K and *Pf*PI4K, we generated linear plasmids capable of integrating the LacI-*lacO* conditional regulation system into the native *Pf*Kelch13 5’ UTR locus. We designed versions of the construct both with and without HA tags at the N-terminus but were not able to recover N-terminally tagged parasites after two transfection attempts. We successfully recovered transfected NF54::pCRISPR parasites^13^ with an untagged *Pf*Kelch13 - regulating construct, transfected in the presence of IPTG. Upon removal of IPTG, parasites showed stalled growth for two IDCs before dying (Fig. 5A). Microscopy showed parasites were unable to progress to trophozoite life stages in the second IDC following IPTG removal (Fig. 5B), as observed in previous work using conditional knockout of *Pf*Kelch13^8^. We validated *Pf*Kelch13 knockdown through qPCR 72 h after removal of IPTG (Fig. 5C). We also confirmed no interaction was detectable between *Pf*Kelch13 and DHA using constant drug, constant expression IC_50_ shift assays (Fig. 5D, Fig. S12), as found in other studies^67–69^. No interactions were observed with other drugs used in this study.

**Figure 5.**
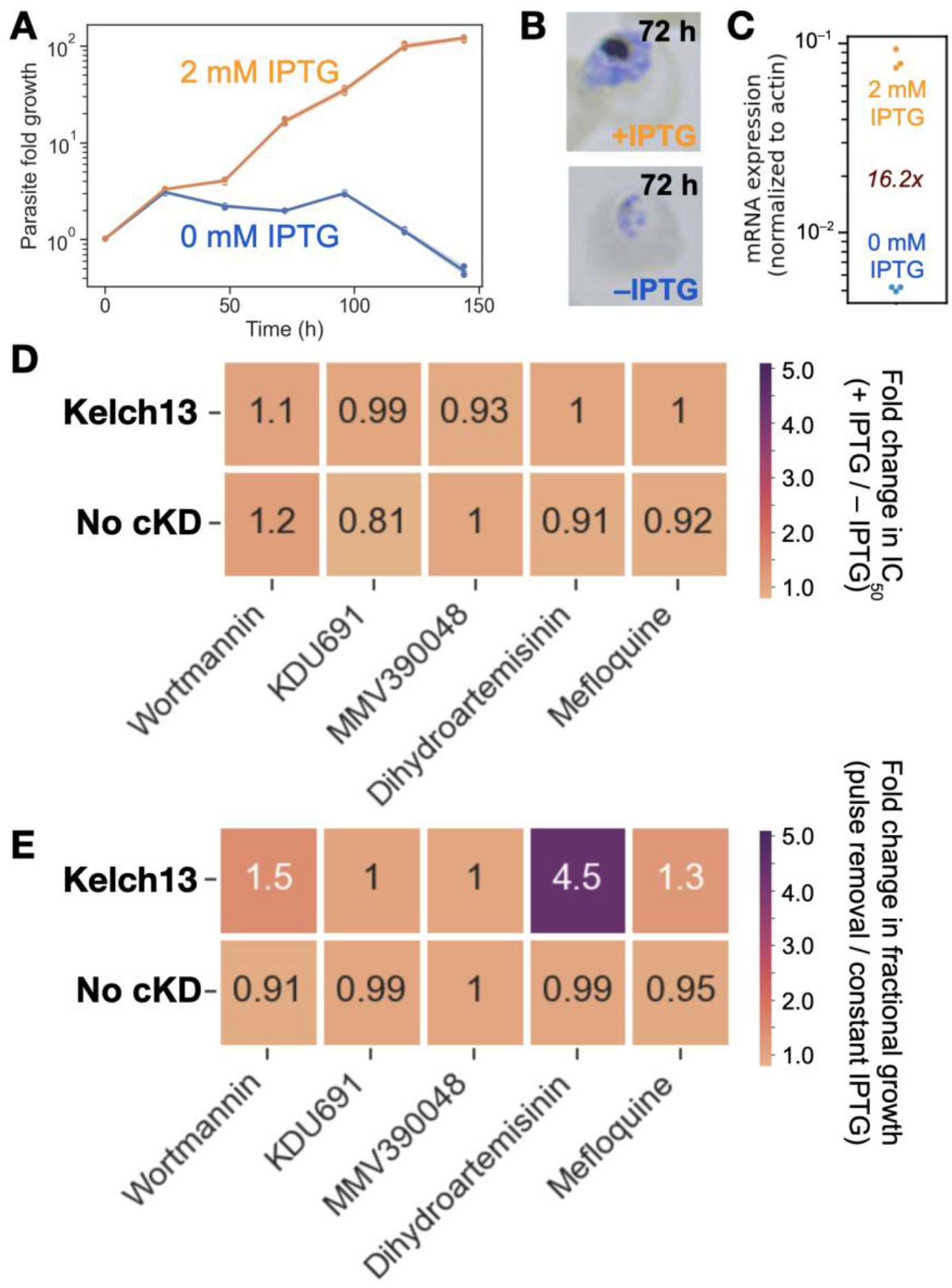
LacI-based transcriptional knockdown shows essentiality and drug interactions of *Pf*Kelch13. Parasite growth following knockdown of *Pf*Kelch13 by removal of the small molecule inducer IPTG. Parasite morphology after 72 h of growth with or without IPTG for *Pf*Kelch13 knockdown. **(C)** RT-qPCR measurement of *Pf*Kelch13 mRNA abundance (normalized to actin) after 72 h of growth with and without IPTG. **(D)** Fold change in IC_50_ of antimalarial compounds, including known *Pf*Kelch13 inhibitor dihydroartemisinin (DHA), during transcriptional knockdown of *Pf*Kelch13. **(E)** Fractional survival of parasites in the next intraerythrocytic development cycle (IDC) after exposure to a 6 h pulse of 700 nM DHA following temporary knockdown of *Pf*Kelch13 expression.

We next inquired whether temporary removal of *Pf*Kelch13 could rescue parasites from antimalarial exposure. We exposed parasites to a 6 h pulse of an antimalarial compound, testing DHA as well as the compounds wortmannin, KDU691, MMV390048, and mefloquine. We inquired whether temporary removal of *Pf*Kelch13 expression 24 h before compound exposure resulted in changes to parasite survival, measured one IDC after the exposure pulse and the restoration of *Pf*Kelch13 expression by adding IPTG. We compared growth with and without temporary removal of *Pf*Kelch13 expression in the presence of different compounds as a fraction of total growth in the same *Pf*Kelch13 expression condition with no compound. The results suggest a strong increase in parasite survival and growth in the presence of DHA when *Pf*Kelch13 expression was temporarily removed (Fig. 5E). The fact that temporary knockdown of *Pf*Kelch13 expression and DHA exposure can detect a difference in survival outcome underscores the potential of LacI-based transcriptional knockdown as a tool to carry out compound-gene interaction studies.

## Discussion

Robust regulation of transcription has long proven to be a challenge in malaria parasites. In this work, we describe how inducible control of gene transcription can be achieved in *P. falciparum* using synthetic operator arrays and their cognate bacterial repressor proteins. We present two different transcriptional control systems built using the bacterial repressors TetR and LacI, inducible by two different inducer molecules. The TetR-based system achieves 15- to 50-fold regulation in trophozoites and schizonts, improving on existing methods of inducible transcriptional control by an order of magnitude^21^. The LacI-based system is able to achieve 200- to 800-fold regulation of expression, outperforming not only previous methods of transcriptional control, but all previous conditional knockdown systems available in *Plasmodium* parasites, the most effective of which is currently the TetR-DOZI aptamer system with around 100-fold regulation of translation^11,13^. Using LacI-based modulation of transcription, we validated two lipid kinases that had previously escaped effective knockdown as essential genes and confirmed them to be specific targets of small molecule inhibitors using IC50 shift assays. This demonstra tes how stronger knockdown techniques can elucidate gene essentiality and gene-compound interactions that other technologies might miss. Additionally, we conditionally regulated *Pf*Kelch13 and took advantage of the reversibility of this conditional knockdown system to study the effects of temporary removal of expression on artemisinin survival. The increased survival of *Pf*Kelch13 knockdown parasites shows that temporary knockdown tools such as LacI-based transcriptional control could be used to assay or screen for compound-gene interactions not visible in other assays. In all, these results demonstrate the flexibility and robustness of synthetic transcriptional regulation in *P. falciparum* for functional genomics and parasite engineering.

Despite the successes and potential of these transcriptional regulation systems, it is important to consider their limitations. The pPC101 plasmid system applied to native genes in this study replaces a target gene’s native promoter. This can affect normal transcriptional dynamics, as seen with the reduction in *Pf*PI4K expression. Nevertheless, we have seen here how other knockdown techniques can also affect native expression patterns, and the example of *Pf*PI4K demonstrates that valuable information and assays can still be conducted even when the native expression regime of a gene of interest is altered, provided the dynamic range of the system is great enough. If editing the 5’ extreme of a gene were impossible for a given application, integration of *lacO* arrays at the 3’ end still achieved measurable change in expression.

The toxicity of NLS-tagged TetR to parasites remains an open question. TetR binding to arrays of *tetO* sequences has been known to cause a steric roadblock to the DNA replication complex leading to toxicity in prokaryotes^74^. This would explain why toxicity in our studies was only observed when both the protein and DNA components of the nuclear-localized TetR-*tetO* interaction were present, and why toxicity was relieved upon breaking this interaction with aTc. However, it is unclear why this was not observed with the interaction between nuclear-localized LacI and its own cognate operator, *lacO*, particularly given the stronger transcriptional repression that LacI-based systems were found to have here.

Indeed, another outstanding question is why LacI-based regulation showed such a larger dynamic range than TetR did. LacI is reported to have 4-fold higher affinity for its DNA operator than TetR *in vitro*^75–77^. Additionally, LacI is a larger protein than TetR and forms tetramers, not dimers, when bound to DNA, possibly increasing steric hindrance to the pol II transcription complex. However, this does not explain why studies in mammalian systems have found TetR-based systems to achieve a dynamic range more than 10-fold greater than that of LacI^42–45^. It would seem that the mechanisms determining the effectiveness of even simple recombinant tools built around steric hindrance can be species-specific — an important reminder for synthetic biology at large.

Regardless, the tools described in this work represent powerful alternatives for conditional expression in *P. falciparum* and open new avenues for research in malaria parasites. The transcriptional nature of these tools makes them amenable to exploring noncoding RNA biology, such as the regulation of *var* gene expression by long, noncoding RNA (lncRNA)^32,41^. The orthogonal nature of IPTG-LacI-*lacO* and aTc-TetR-*tetO* (including TetR-DOZI) systems can also allow for studies of interactions between two independently regulated genes. These tools open the possibility for engineering tunable synthetic gene circuits in malaria parasites, with applications in metabolite biosensing, immunoengineering, and drug discovery.

## Acknowledgements

We are grateful to Narendra Maheshri for providing the original 180-repeat *tetO* plasmid used as a PCR template for DNA assembly, Stephen Goldfless for the original pSG372 plasmid used as a testbed, and Suresh J. Ganesan for the pINT plasmid without neomycin resistance (pSG234). We also thank Cassandra Rogers and the Whitehead Keck Microscopy Core for training and assistance with fluorescence microscopy, Bogdan Fedeles of the MIT Center for Environmental and Health Sciences and the staff of the Broad Institute Flow Cytometry Core for training and maintenance of cytometry facilities, and the staff of the MIT and Koch Institute BioMicro Center for training and maintenance of qPCR equipment. We deeply thank Kyle McLean, Lisl Escherick, Gaël Chambonnier, Charisse Pasaje, Luiz Godoy, Khan Osman, Leah Imlay, and other current and former members of the Niles lab for their mentorship and guidance throughout this project. P.C. received support from the Surpina and Panos Eurnekian Biotechnology Fund through the MIT Office of Graduate Education and the Siebel Scholars Program. This work was supported by grants from the Bill and Melinda Gates Foundation (OPP1162467 and OPP1158199), Broad Next10, National Institute of General Medical Sciences Center for Integrative Synthetic Biology Grant (P50-GM098792), National Institutes of Environmental Health Sciences Core Center Grant (P30-ES002109).

## Data and code availability

All plasmid maps, data, and analysis code is deposited in a GitHub repository archived with a digital object identifier (DOI) at doi.org/10.5281/zenodo.12193161. Code for GeneTargeter can be found at github.com/pablocarderam/genetargeter (archived under DOI zenodo.org/doi/10.5281/zenodo.12193163), and the software is available for use at genetargeter.mit.edu.

## Author contributions

P.C. and J.C.N. designed all experiments in the study. S.S. helped design and carry out experiments for compound-target interaction assays. S.D. created conditional knockdown lines for PI3K and PI4K using the TetR-DOZI system. P.C. conducted all experiments and data analysis in the study, as well as code development for GeneTargeter. P.C. drafted the text, and all authors read and edited the manuscript.

