## Supplementary methods, figures, and tables for "Synthetic transcriptional control in the malaria parasite *Plasmodium falciparum*"

#### *General DNA assembly*

Two different plasmid systems were used throughout this study. The first is based on pSG372[1] and consists of two diverging transcriptional units with back-to-back *P. falciparum* promoters mounted on a circular plasmid containing *attP* sites used for BxbI-mediated integration into *attB* loci on transgenic parasites[2]. This plasmid system was used to assemble constructs used to test regulation of gene expression by modifying the repressor proteins being used for regulation, their cognate operator sequences, the output proteins to be regulated and measured through luminescence or fluorescence, and the promoters driving the expression of these proteins. The second type of plasmid used was a linear plasmid backbone based on the pJAZZ system (Biosearch Technologies) and following the structure of plasmid pSN150[1]. These linear plasmids were used to assemble repair templates for native gene modification through Cas9-mediated homologous recombination.

To assemble these different constructs, plasmid backbones were digested with restriction enzymes and new inserts were obtained through polymerase chain reaction (PCR) amplification, DNA synthesis, or oligonucleotide annealing and extension, according to the type of assembly. Bacterial cultures containing plasmids were grown overnight in Terrific Broth with 200 rpm orbital shaking at 37 $^{o}$C (unless otherwise noted, see Supplementary Table S1). DNA extraction through minipreps and midipreps was carried out according to QIAGEN manufacturer protocol and eluted in Milli-Q (Sigma Aldritch) purified water. Restriction enzyme digestions were carried out according to New England Biolabs manufacturer protocol in 40 μL reactions and incubated for 6 h before inactivation. Digestion products were purified directly from reaction using QIAGEN DNA Purification and Clean-up kits. PCR reactions were carried out using Kapa HiFi 2X Master Mix (Sigma Aldritch) in 50 μL reactions with 5% DMSO by volume and a PCR protocol designed according to manufacturer's specification, unless otherwise noted below. Minipreps were used as templates whenever possible, but for amplification of *P. falciparum* DNA, infected blood was used as a template (2 μL of a solution composed of 40 μL of water and approximately 0.2 μL of infected blood with culture media removed). PCR products were run in 4% agarose gels at 120 V for 20 minutes and visualized with ethidium bromide. Amplicon bands were cut and purified using QIAGEN DNA Purification and Clean-up kits. Plasmids were assembled using Gibson Assembly[3] with Telesis Bio Gibson Assembly Ultra Kits according to manufacturer specifications. A single insert was inserted into a backbone (linear backbones are constituted of two separate fragments) at every step to improve efficiency, necessary for some difficult assemblies involving AT-rich *P. falciparum* DNA or highly repetitive operator sequences. Gibson Assembly products were dialyzed on membranes and transformed by electroporation using a BioRad electroporator with default "Ec1" settings. All circular plasmids were transformed and grown with kanamycin in New England Biolabs 10-beta electrocompetent cells, while linear plasmids were transformed into BigEasy v2 cells (Biosearch Technologies, custom order) and grown with chloramphenicol. Cells were recovered in Terrific Broth for ~40 min before being plated on TB agar with the appropriate antibiotic selection marker grown overnight. Colonies were picked with wooden toothpicks, grown overnight in liquid culture, and Sanger sequenced (Azenta Life Sciences). Linear whole-plasmid sequencing (Plasmidsaurus Inc., using Oxford Nanopore Technology) was carried out on finished constructs. Bacterial cultures containing successful constructs were stored at –80 $^{o}$C in 33% glycerol by volume.

Details for each type of assembly are provided below and descriptions of all plasmids, oligonucleotide sequences, and links to DOI-archived plasmid map GenBank files may be found in Supplementary Table S1.

*tetO operator copy number variant assembly*
Plasmid pPC000 was prepared by removing the TetR aptamer from pSG372 using AflII and XhoI and resealing the plasmid with annealed oligos. Arrays of *tetO* sequences of different lengths were obtained by PCR amplification from plasmid pRS316 (courtesy of Dr. Narendra Mahesri) and inserted into pPC372 using AflII to obtain plasmids pPC001–pPC005 or using both AflII and XhoI to obtain plasmids pPC006–pPC010.

*Repressor and operator variant assembly*
Plasmid pPC011 was built by replacing a transcriptional unit in pPC000 (digested by AgeI and AvrII) with PCR products containing modified overhangs to insert an XmaI restriction enzyme cut site downstream of the TetR sequence in plasmid pPC000, adding a modular SV40 nuclear localization signal at the start of the TetR sequence, and replacing firefly luciferase with *Cypridina* luciferase (obtained via Integrated DNA Technologies gBlock DNA synthesis, but ultimately found not to function in *P. falciparum*) on the opposite transcriptional unit. Firefly luciferase ultimately replaced *Cypridina* luciferase once again through PCR amplification of the insert and digestion of the backbone with XhoI and SpeI.

We chose to test bacterial repressor systems that are commonly used in synthetic biology applications, including CymR-*cuO*, inducible by cumate; PhlF-*phlO*, inducible by 2,4-diacetylphloroglucinol (DAPG), and LacI-*lacO*, inducible by lactose and analogues such as isopropyl β-D-1-thiogalactopyranoside (IPTG). Because the operator sequence for PhlF contained a positive strand ATG sequence, the sequence was altered to avoid a premature start of translation before the start of the coding sequence. All repressors were fused to SV40 NLS and all operator sequences were spaced by 2–3 bp. Bacterial repressor sequences were synthesized as gBlock gene fragments (Integrated DNA Technologies). These gene fragments were inserted into pPC030, replacing the TetR sequence excised with BamHI and XmaI. Arrays of operators corresponding to each repressor being used were synthesized by overlap extension PCR using 200 bp Ultramer oligonucleotides (Integrated DNA Technologies) due to the high repetitiveness of these sequences causing difficulties for standard gene fragment synthesis. These arrays were then inserted into each corresponding plasmid by digesting with AflII, generating plasmids pPC032–pPC042 and pPC088–pPC089. The operator array in plasmid pPC038 was found to be unstable when grown at 37 $^{o}$C and was therefore grown at 28 $^{o}$C instead. Plasmid pPC070 was constructed by excising the SV40 nuclear localization sequence in pPC032 using BamHI and AvrII. Plasmid pPC126 was assembled by removing the *lacO* mixed array from plasmid pPC089 through excision by AflII and XhoI, followed by reinsertion of the operator array (obtained through PCR amplification) downstream of the FLuc coding sequence by digestion with SpeI.

*Low-expression TetR variant assembly*
Plasmid pPC071 was constructed by replacing the HSP86 promoter in pPC032 (excised using AscI and AvrII) with a mRPL2 promoter, obtained by PCR. PCR of *P. falciparum* promoter sequences presents a significant challenge, as these noncoding sequences commonly exceed 90% AT richness. To successfully PCR promoters, long primer annealing sequences and overhangs were used. PCR protocols were modified to have five "sub-cycles" within each annealing step in which the temperature was alternated between the chosen annealing temperature (commonly 58 °C) and a higher "pseudo-extension" temperature (commonly 63 °C). Extension temperatures were lowered to 65 °C from 72 °C if no amplification was observed, with an increased extension time (commonly 5 minutes per cycle). Plasmid pPC072 was assembled by adding a Sir2A sequence (also obtained through PCR) at the end of the TetR sequence with a 10-residue glycine-serine linker within the XmaI cut site.

*Output promoter variant assembly*
Promoter sequences were obtained via PCR using the "sub-cycling" method described above and inserted into plasmid backbones derived from AscI and AflII digestions of either pPC071 or pPC089, depending on the repressor system to be used. These assemblies yielded plasmids pPC094–pPC097 and pPC111–pPC114, respectively.

*Output gene variant assembly*
Plasmids pPC071 and pPC089 were digested with XhoI and SpeI to replace the firefly luciferase coding sequence with that of fluorescent protein mNeonGreen, synthesized as a gBlock gene fragment from Integrated DNA Technologies. This yielded plasmids pPC098 and pPC099, respectively.

*Native gene conditional expression construct assembly*
Plasmid pPC101 was obtained by amplifying the contents of plasmid pPC089 (comprising the expression cassette encoding for SV40_LacI, *Renilla* luciferase, and blasticidin S-deaminase as well as the regulated CAM promoter with *lacO* operator array) and inserting the product into a linear pJAZZ backbone obtained by digesting plasmid pSN150 with FseI and AsiSI. This plasmid constitutes the base system used to build conditional expression constructs. To build expression constructs regulating PI3K, PI4K, and Kelch13, sgRNA, left and right homology regions, and recoded region sequences were obtained from GeneTargeter (genetargeter.mit.edu) using the pPC101 framework setting. PCR primers for the left homology region were extended to improve amplification efficiency, and PCRs were carried out using the "sub-cycling" method described above. The PCR products were inserted into pPC101 by digesting with FseI. A single gene synthesis fragment was designed to comprise optional HA tags, the recoded region, the right homology region, and the sgRNA expression cassette. These fragments were synthesized by Twist Biosciences or as gBlocks from Integrated DNA Technologies according to synthesis difficulty and integrated into the corresponding linear plasmids digested with AsiSI.

*Parasite culture*

*P. falciparum* 3D7 NF54 parasites were cultured at 37 °C, 5% $O_2$, 5% $CO_2$, and 2% hematocrit in RPMI complete media (RPMIc), prepared using 10.8 g/L RPMI-1640 (US Biological), 2.5 g/L Albumax II (Life Technologies), 2 g/L sodium bicarbonate, 25 mM HEPES pH 7.4 (pH adjusted with potassium hydroxide), 1 mM hypoxanthine and 50 mg/L gentamicin. Human red blood cells (purchased from Research Blood Components, Watertown, MA) no more than two weeks old were used for parasite cell culture. Plasma was removed and red blood cells were washed three times with equal volume of RPMI wash media (10.8 g/L RPMI-1640, 25 mM HEPES, and 50 mg/L gentamicin) by centrifugation at 2000 rcf for 5 min. Red blood cells were stored at 50% hematocrit. Parasites expressing Cas9 and T7 (parasite line NF54::pCRISPR[1]) were cultured in 2.5 nM WR99210 (Jacobus Pharmaceuticals). Regular culture was carried out in 10 mL parasite cultures inside vented culture flasks with media changes every other day. To prepare smears, 5 µL of infected red blood cells were smeared on glass slides, air dried, fixed for 30 s in methanol, stained for 10 minutes in 10% volume modified Giemsa stain (Sigma Aldritch), washed in water and air dried. Parasitemia was assessed via light microscopy with an 100X oil objective and cultures were diluted periodically to keep parasitemia below 10%.

*Parasite storage and recovery*

Parasites were stored by mixing 200 µL of 5–10% parasitemia infected red blood cell pellets (centrifuged at 450 rcf for 5 minutes) with equal volume of glycerolyte 57 (Fresenius Kabi) and immediately freezing at –80 °C. Frozen cultures were recovered by thawing vials and adding a series of thawing solutions. First, 65 µL of 12% weight/volume sodium chloride were added, followed by 6.5 mL of 1.6% w/v sodium chloride added slowly (1 drop/s). Cultures were centrifuged (450 rcf, 5 min) and supernatant was removed. A third solution containing 0.9% w/v sodium chloride and 2% w/v glucose was added, followed again by centrifugation and supernatant removal. Finally, infected red blood cell pellets were washed in RPMIc before being transferred into a culture flask.

*Parasite transfection*

Transfections were performed using a modified version of the red blood cell preloading method[4]. DNA for transfection was obtained by midiprep (QIAGEN Plasmid Midi Kit). Approximately 100 µg of DNA (at concentrations of 200–1000 ng/µL) were added to 300 µL of freshly-washed and pelleted red blood cells no more than a week old. All circular plasmid transfections (all transfections in this study except for conditional knockdowns of native *P. falciparum* genes) contained two *attP* recombination sites, enabling genomic integration in *P. falciparum* parasites with an *attB* recombination site inserted into the *cg6* (PF3D7_0304500) locus when co-transfected with plasmid pSG234 expressing Bxb1 integrase[2]. In these cases of cotransfection with pSG234, similar molar amounts of each plasmid being transfected were used. Red blood cells were electroporated in 0.2 cm cuvettes (8 square wave electroporation pulses of 365 V, 1 ms/pulse, 0.1 ms intervals). Electroporated cells were then washed four times with 1 mL RPMIc (centrifugating at 2000 rcf for 2.5 minutes) and added to 10 mL of RPMIc. These cultures were then infected with 1 mL of parasite cultures at 5% parasitemia. Media was changed daily for the first five days after transfection, and selection was started on the fourth day using 2.5 µg/mL blasticidin (RPI Corp, B12150-0.1). 150 µL of 50% hematocrit were added to cultures on the day selection began. For transfection of essential gene knockdown constructs such as PI3K, PI4K, and Kelch13 knockdowns, 2 mM IPTG inducer was used in culture media from the moment of transfection. Parasites were monitored for up to five weeks after transfection by smears and weekly *Renilla* luciferase measurements obtained from blood pellets from 200 µL culture samples assayed with Promega *Renilla*-Glo(R) Luciferase Assay System on a GloMax 20/20 luminometer (Promega). Transfected cultures were frozen for storage as soon as possible.

*Gene regulation assays*

To assay regulation of gene expression, 4 mL sorbitol-synchronized[5] cultures were grown in triplicate on 12-well plates at each concentration of inducer (aTc or IPTG) as indicated. Inducer was removed from culture media by washing and centrifugating (5 min, 450 rcf) cultures three times in equal volume of media with no inducer. Cultures were sampled daily. Two 200 μL culture samples were taken and centrifuged at 5000 rcf for 1 minute. Cell pellets were stored at –80 °C. *Renilla* luciferase (RLuc) was measured using *Renilla*-Glo(R) Luciferase Assay System (Promega) on a Glomax 20/20 luminometer (Promega). If relevant to the experiment, firefly luciferase (FLuc) was assayed with Promega ONE-Glow Luciferase Assay System (Promega). To quantify the magnitude of FLuc regulation, FLuc luminescence was normalized to constitutive RLuc luminescence. This normalized expression was then calculated as a fraction of the maximum normalized expression of an equivalent genetic construct containing all the same promoters, terminators, and coding sequences, but with no operators affecting FLuc expression.

*Flow cytometry and fluorescence microscopy*

To assay regulation of gene expression, 4 mL sorbitol-synchronized[5] cultures were grown in triplicate on 12-well plates at each concentration of inducer (aTc or IPTG) as indicated. 500 μL samples were stained with 0.1 μL Syto61 (ThermoFisher) for 30 minutes and filtered. Fluorescence was measured on a CytoFLEX flow cytometer (Beckman). Data processing was carried out with Cytoflow[6]. Fluorescent microscopy was carried out using a DeltaVision II (Image Solutions).

*Compound growth inhibition assays*

To assay drug-target interactions, 200 μL sorbitol-synchronized[5] cultures were grown in triplicate on 96-well plates at each concentration of drug and inducer as indicated. Inducer was removed from culture media by washing and centrifugating (5 minutes, 450 rcf) cultures three times in equal volume of media with no inducer. After 72 h, 50 μL of supernatant was removed from each well and *Renilla* luminescence was assayed using *Renilla*-Glo(R) Luciferase Assay System (Promega) on a GloMax Navigator plate reader (Promega).

*Compound pulse assays*

Compound pulse assays were carried out in triplicate 1 mL cultures. At 24 h after synchronization, parasites were exposed to approximately 100 times the $IC_{50}$ value for each compound (except for wortmannin, which has an exceptionally high $IC_{50}$): 200 μM wortmannin (about 25 times $IC_{50}$), 2.5 μM KDU691, 1.5 μM MMV390048, 700 nM DHA, and 1 μM mefloquine. At 30 h after IPTG removal, cultures were washed three times to remove DHA. We decided to apply the compound pulse to parasites during the trophozoite stage instead of the ring stage, as commonly done in ring stage survival assays for Kelch13 resistance, since we replaced the wild-type Kelch13 promoter, which has an expression peak during ring stages, with the calmodulin promoter, which peaks expression during trophozoite stages. IPTG was added back to all cultures after compound pulse exposure, and luminescence was measured following the procedure of *Renilla* luciferase luminescence assays used for gene regulation measurement as outlined above.

*Real-time quantitative PCR*

To assay regulation of gene expression, 4 mL sorbitol-synchronized[5] cultures were grown in triplicate on 12-well plates at each concentration of IPTG as indicated. During sample collection, RNA was isolated from the entire culture using RNEasy RNA extraction kits (QIAGEN). RNA was used as template in 20 µL reverse transcriptase and PCR combined reactions using the NEB Luna kit (New England Biolabs) in 96-well plates. Each biological replicate was measured in three technical replicate reactions. RT-qPCR measurements were obtained using a LightCycler 480 II Real-time PCR Machine (Roche). Measurements of each gene being probed were normalized to actin as an endogenous control, using primers obtained from other work[7]. All primers used are available in Supplementary Table S2.


### Supplementary Figures

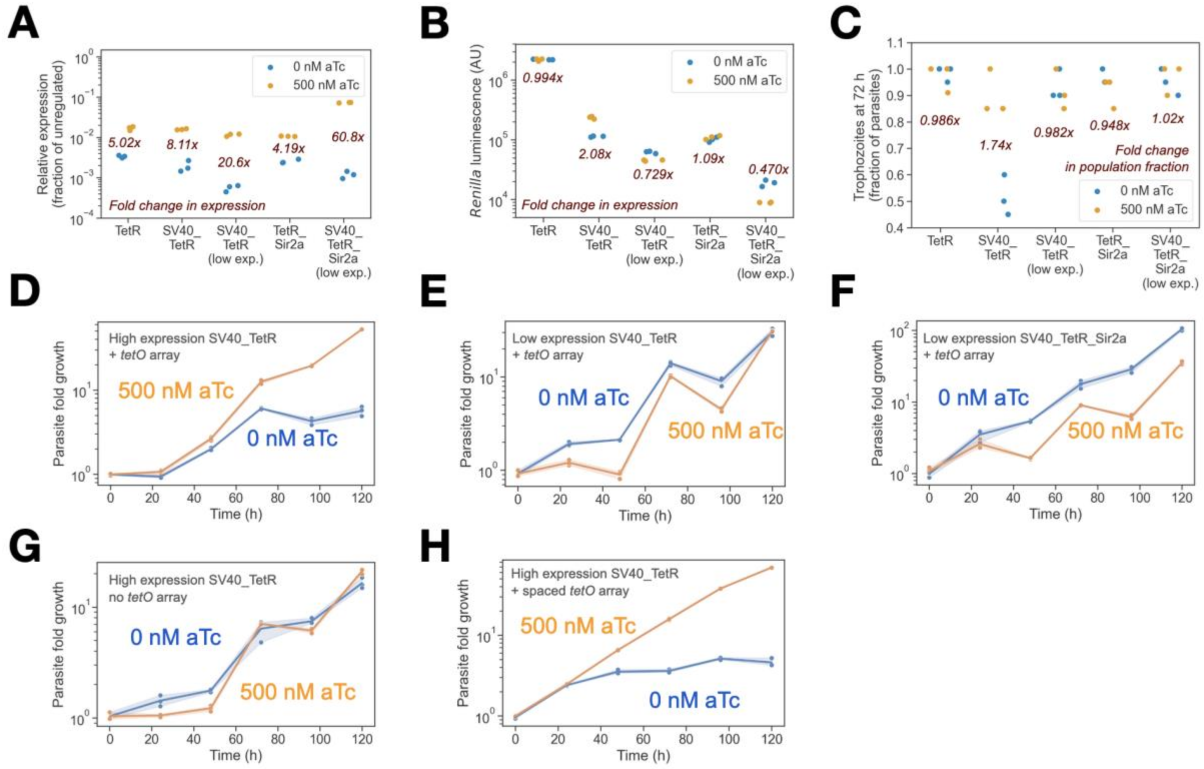


**Figure S1.** Rational engineering of an effective, nontoxic transcriptional regulator. **(A)** Addition of an SV40 nuclear localization signal to the TetR repressor results in increased fold change in the regulation of the target gene, firefly luciferase (FLuc). Fusion of an Sir2A histone deacetylase protein to TetR appears to further increase fold regulation, but **(B)** is in fact an artifact of increased off-target expression of *Renilla* luciferase from the adjacent transcriptional unit when in the absence of anhydrotetracycline (aTc). **(C)** Low expression of SV40-fused TetR from a weak promoter is required to avoid the toxic arrest of parasites at the ring stage. **(D)** With high expression of SV40, parasite growth decreases and then stops after 72 h in the absence of aTc inducer. With a nontoxic, low expression system for **(E)** SV40-fused TetR and **(F)** SV40- and Sir2a-fused TetR, parasites can grow freely without inducer aTc. **(G)** High-expression SV40-fused TetR does not inhibit growth when *tetO* sequences are not present. **(H)** Using the original *tetO* array with longer spacers results in the same toxic inhibition of growth when in absence of aTc.

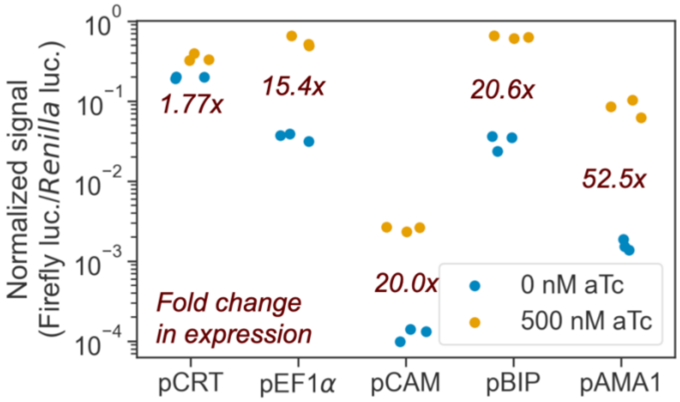


**Figure S2.** Robust regulation is observed when applying the TetR-based system on multiple parasite 5’ UTRs with promoters active in trophozoite, schizont, and merozoite stages, but is not as effective when regulating a 5’ UTR for chloroquine resistance transporter (CRT), which contains a promoter active in the ring stage. In each case, an engineered array of eight *tetO*-based sequences is inserted between the 5’ UTR and the gene being regulated, which here is firefly luciferase (FLuc).

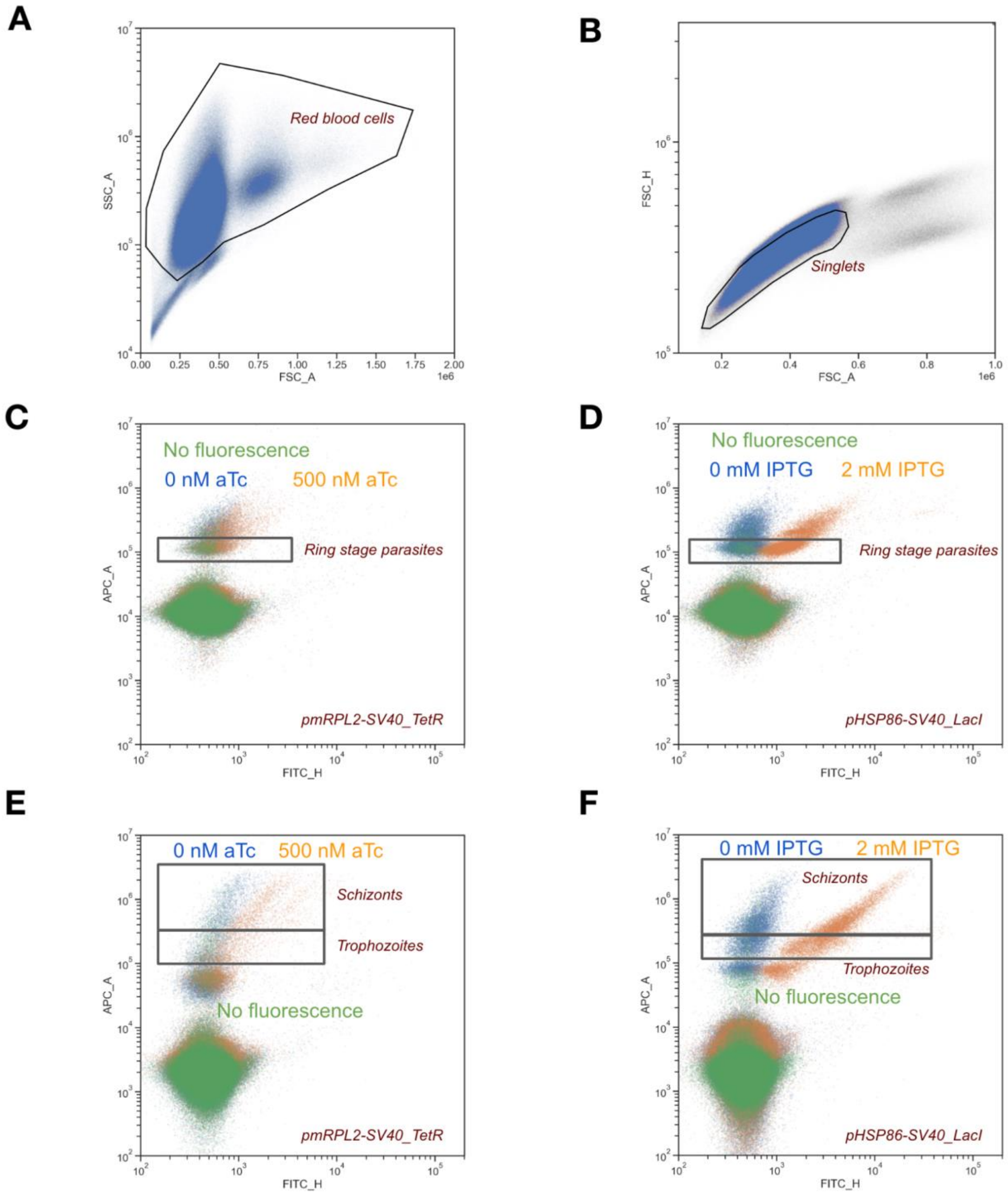


**Figure S3.** Gating scheme used to characterize synthetic transcriptional regulation of mNeonGreen. **(A)** Red blood cells (RBCs) and **(B)** singlets were gated and analyzed for red (allophycocyanin channel, APC) fluorescence, denoting parasite-infected RBCs stained with nucleic acid dye Syto61, and green fluorescence, indicating induced expression of mNeonGreen. Cells enriched for early (ring) life stages for **(C)** the TetR-based system and **(D)** the LacI-based system were analyzed in one experiment. A separate experiment focused on cells enriched for late life stages for **(E)** the TetR-based system and **(F)** the LacI-based system.

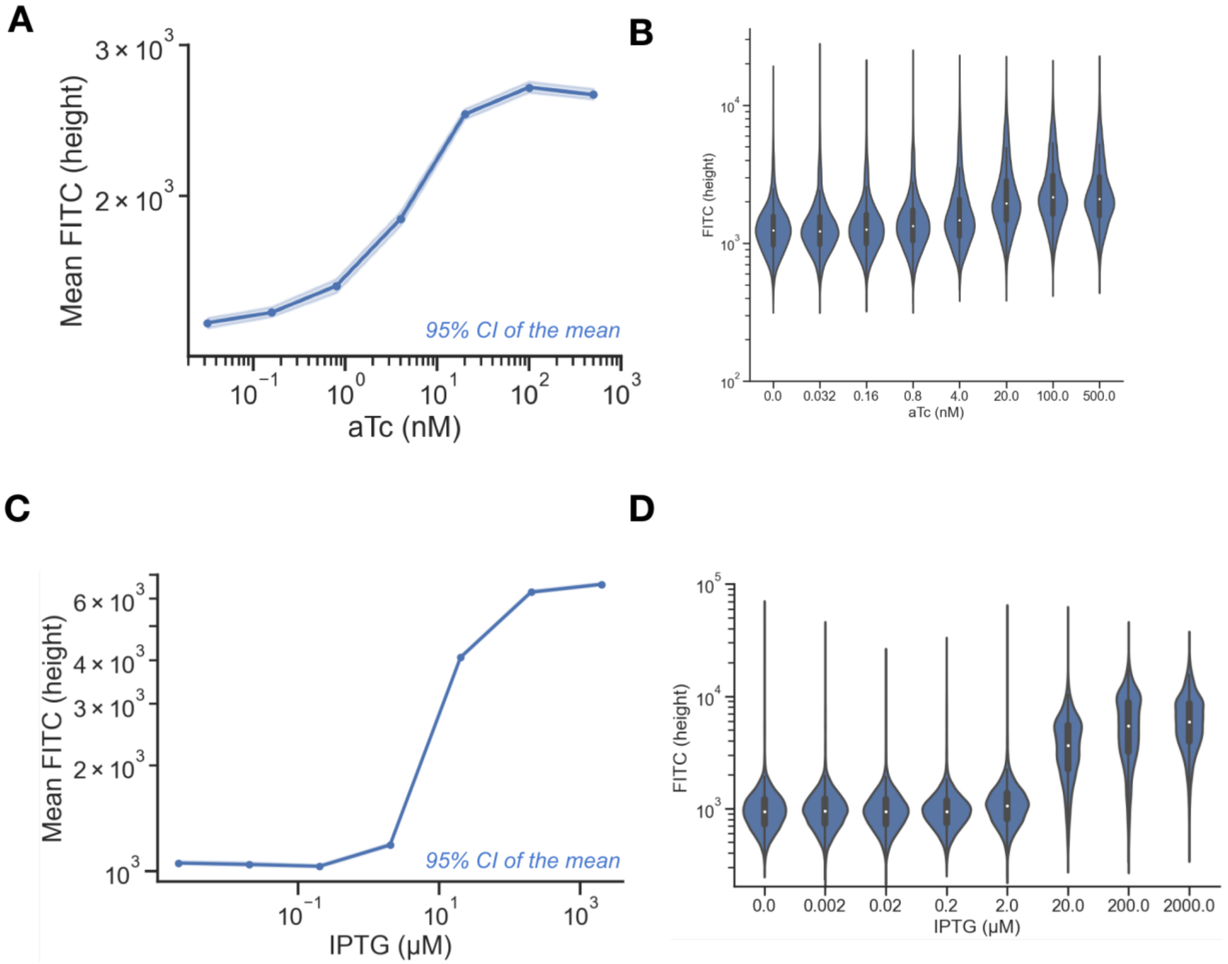


**Figure S4.** Flow cytometry measures the distribution of expression levels in individual cells with synthetic transcriptional regulation. Regulation is seen in both **(A, B)** the TetR-based system and **(C, D)** the LacI-based system. In both cases, regulation of expression in individual cells follows a consistent, monotonic pattern of increase, with no population of cells remaining at a low expression level.

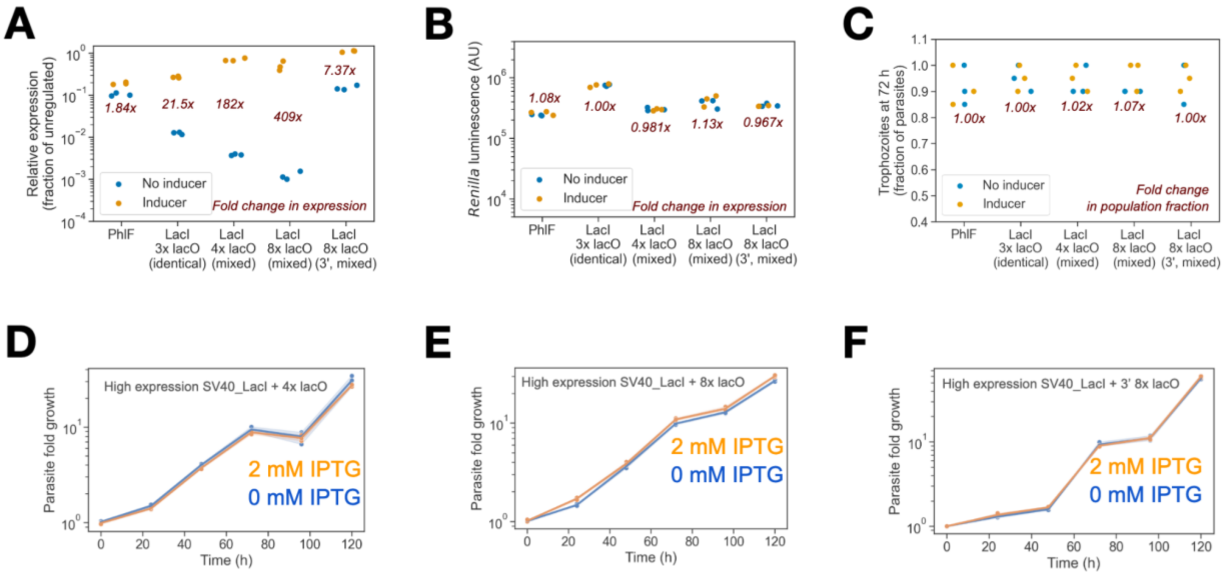


**Figure S5.** Rational engineering of a highly effective *LacI*-based transcriptional regulator. **(A)** PhlF-based, 2,4-diacetylphloroglucinol- (DAPG-) induced regulation was not substantial and the CymR-cuO construct could not be successfully transfected after multiple attempts. Nevertheless, *lacO*-based systems regulated by isopropyl β-D-1-thiogalactopyranoside (IPTG) show considerable regulation. Arrays of mixed variants of *lacO* operators allow for more stable, longer repeats with higher fold regulation of the target gene, firefly luciferase (FLuc), than identical, direct *lacO* repeats. Inserting mixed *lacO* variant repeats at the 3' end of a gene results in regulation to a lesser magnitude than at the 5' extreme. **(B)** No off-target transcriptional effects or differences in growth rate affecting the amount of *Renilla* luciferase luminescence are observed, **(C)** nor are there any effects on parasite life cycle visible by microscopy. Growth rates between induced and uninduced parasites are equal in parasites with **(D)** four 5' *lacO* variant repeats, **(E)** eight 5' *lacO* repeats, and **(F)** eight 3' *lacO* repeats.

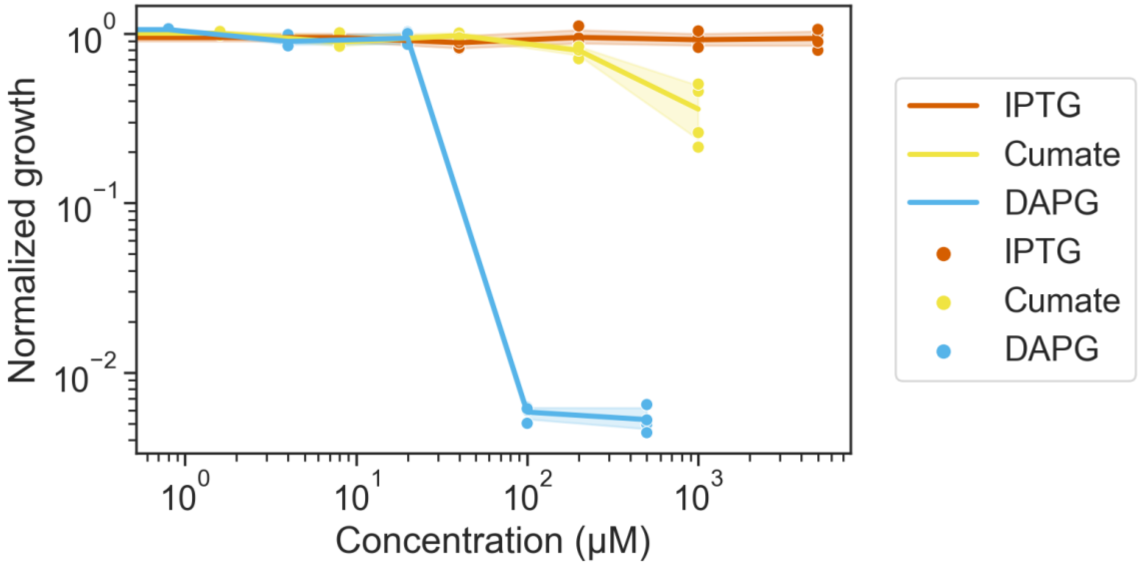


**Figure S6.** Toxicity of inducers for bacterial repressor proteins in *Plasmodium falciparum*. 2,4-Diacetylphloroglucinol (DAPG) and cumate (inducers for the repressors PhlF and CymR, respectively) show parasite growth inhibition at high concentrations. Isopropyl β-D-1-thiogalactopyranoside (IPTG), an inducer of repressor LacI, is found to be nontoxic across all concentrations assayed.

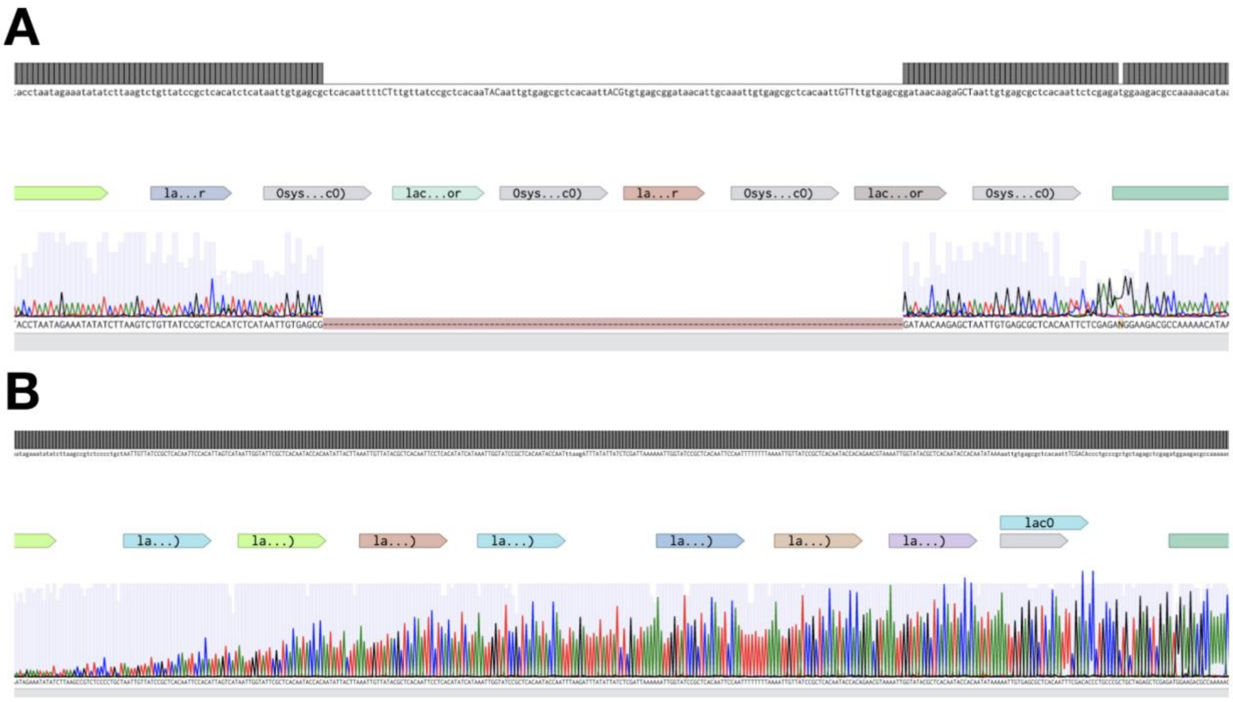


**Figure S7.** Stability of *lacO* array when integrated in parasite genomes. **(A)** Direct *lacO* and *Osys* repeats are unstable above 28 ºC and are lost through recombination during transfection, as shown by Sanger sequencing of the operator array from transfected parasites. **(B)** Mixed *lacO* variant repeats remain stable throughout bacterial growth and parasite transfection at 37 ºC.

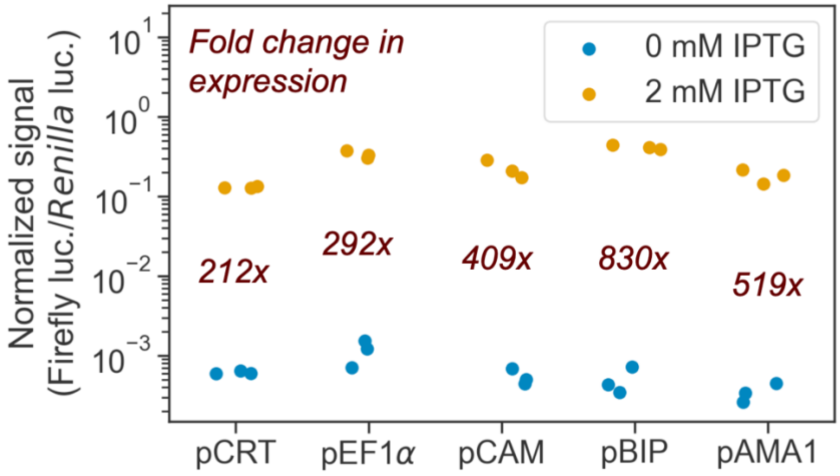


**Figure S8.** Robust regulation of expression is observed when applying the LacI-based regulation on multiple parasite 5' UTRs with promoters active in all red blood cell stages, including rings. In each case, an engineered array of eight *lacO*-based sequences is inserted between the 5' UTR and the gene being regulated, which here is firefly luciferase (FLuc).

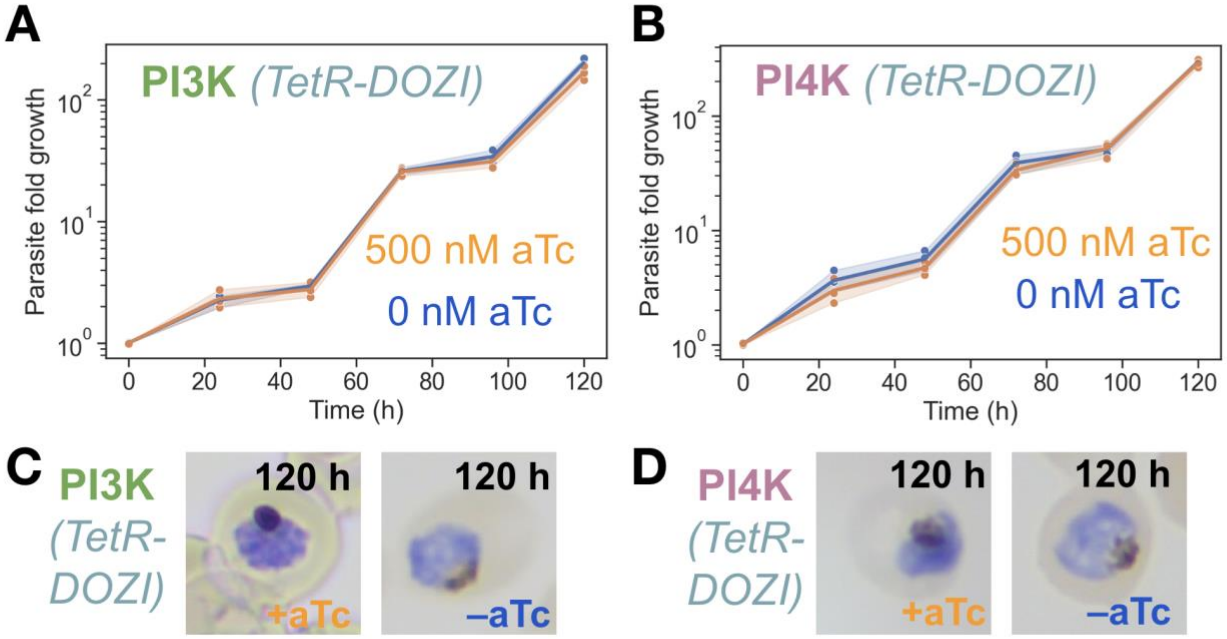


**Figure S9.** TetR-DOZI translational knockdown fails to confirm essentiality of phosphatidylinositol 3-kinase (PI3K, PF3D7_0515300) and phosphatidylinositol 4-kinase beta (PI4K, PF3D7_0509800). *Renilla* luciferase assays show similar patterns of growth for parasites with and without anhydrotetracycline (aTc), which induces translation of the regulated genes, for both (A) PI3K and (B) PI4K. Cell morphology after 120 hours is also similar regardless of aTc inducer presence for both (C) PI3K and (D) PI4K.

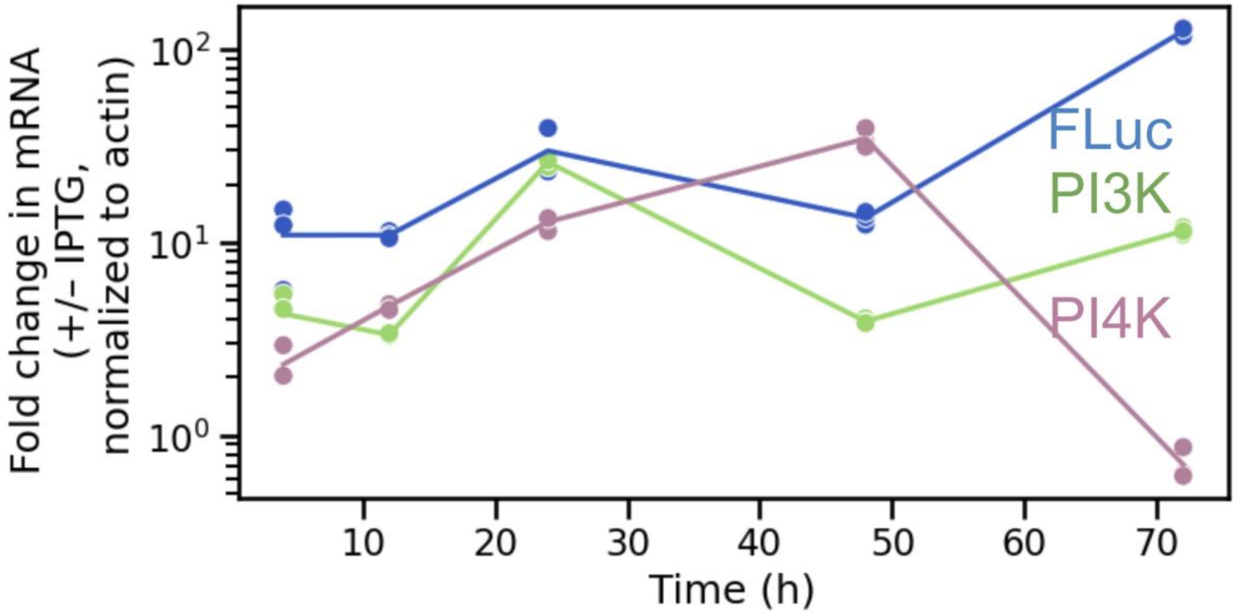


**Figure S10.** RT-qPCR shows performance of LacI-based transcriptional regulation of firefly luciferase (FLuc), phosphatidylinositol 3-kinase (PI3K, PF3D7_0515300), and phosphatidylinositol 4-kinase beta (PI4K, PF3D7_0509800) over time after removal of IPTG from parasite cultures with induced expression of the regulated gene. Values are normalized to actin expression.

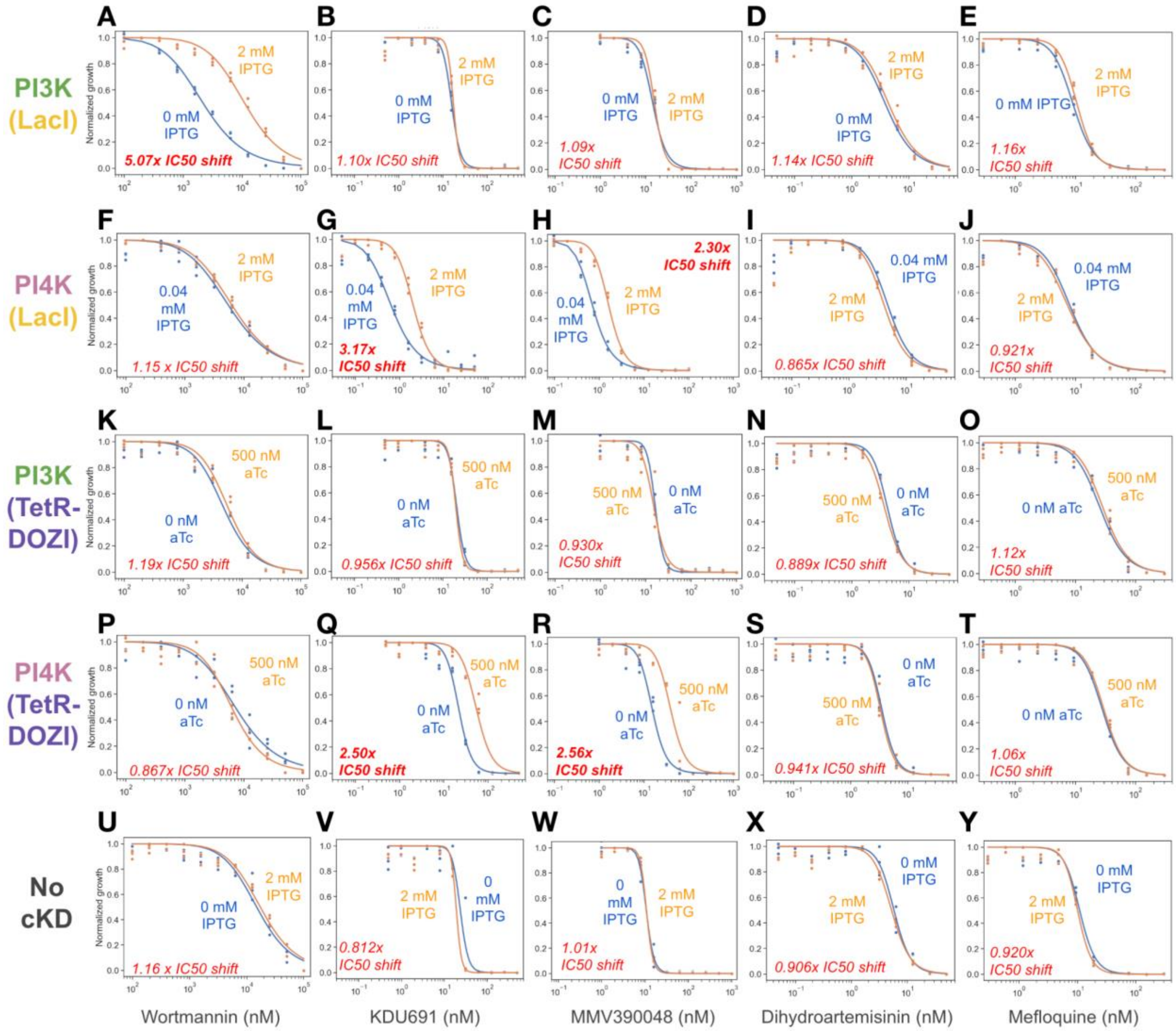


**Figure S11.** LacI-based transcriptional knockdown confirms compound-gene interactions for two *P. falciparum* lipid kinases. **(A)** Phosphatidylinositol 3-kinase (PI3K) is found to be targeted by wortmannin, as evidenced by an isopropyl β-D-1-thiogalactopyranoside- (IPTG-) dependent shift in the half maximal inhibitory concentration (IC50) of the compound when IPTG is removed from the media. **(B–E)** Other drugs show no such IC50 shift. **(F)** Phosphatidylinositol 4-kinase beta (PI4K) is not inhibited by wortmannin but shows an IC50 shift with both **(G)** KDU691 and **(H)** MMV390048. **(I)** DHA and **(J)** mefloquine again shows no IC50 shift with PI4K knockdown. **(K–O)** TetR-DOZI knockdown of expression (inducible by anhydrotetracycline, aTc) does not detect any interactions for PI3K, but **(P–T)** does show the same interaction profile for PI4K. **(U–Y)** In a control line with no LacI expression, no significant IC50 shifts are detected when removing IPTG from growth media, regardless of the compound being studied.

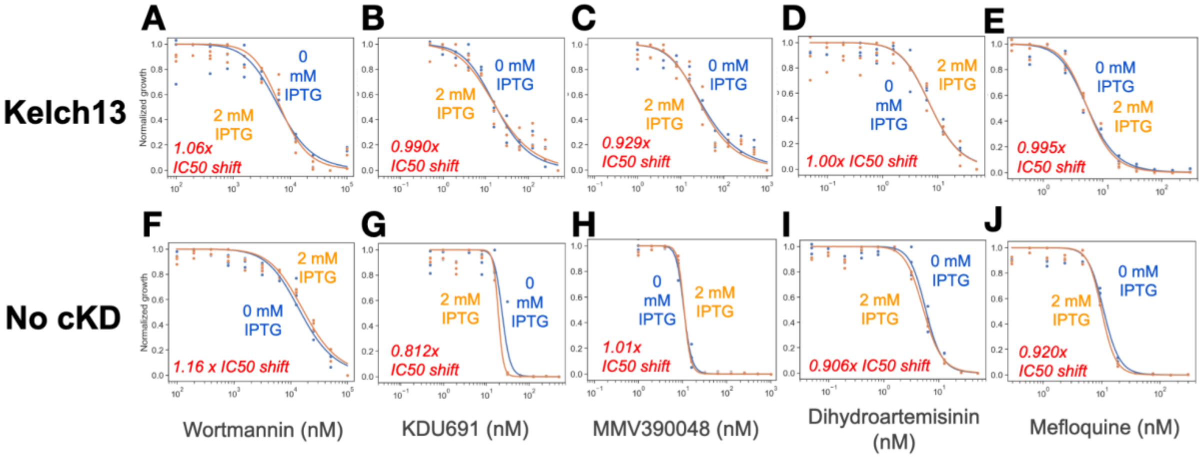


**Figure S12.** LacI-based transcriptional knockdown shows compound-gene interactions for Kelch13. **(A–E)** IPTG-dependent Kelch13 knockdown of expression has no effect on the drug response curve for wortmannin, KDU691, MMV390048, dihydroartemisinin (DHA), or mefloquine. **(F–J)** A control parasite line with no conditional knockdown also shows no differences in $IC_{50}$ regardless of IPTG concentration.

### Supplementary Tables

**Table S1.** Plasmids used in this study.

| Name | Parent Plasmid | Antibiotic | Description |
|---|---|---|---|
| pPC000 | pSG372 | Kanamycin | HSP-Rluc, PfCAM-Fluc, no aptamer. Same as pSG372, but aptamer cut out. |
| pPC002 | pSG372 | Kanamycin | pSG372 with 2x tetO array from Mahesri plasmid downstream of promoter, with 5' aptamer |
| pPC003 | pSG372 | Kanamycin | pSG372 with 4x tetO array from Mahesri plasmid downstream of promoter, with 5' aptamer |
| pPC004 | pSG372 | Kanamycin | pSG372 with 8x tetO array from Mahesri plasmid downstream of promoter, with 5' aptamer |
| pPC005 | pSG372 | Kanamycin | pSG372 with 11x tetO array from Mahesri plasmid downstream of promoter, with 5' aptamer |
| pPC007 | pSG372 | Kanamycin | pSG372 with 2x tetO array from Mahesri plasmid downstream of promoter, without 5' aptamer |
| pPC008 | pSG372 | Kanamycin | pSG372 with 4x tetO array from Mahesri plasmid downstream of promoter, without 5' aptamer |
| pPC009 | pSG372 | Kanamycin | pSG372 with 8x tetO array from Mahesri plasmid downstream of promoter, without 5' aptamer |
| pPC010 | pSG372 | Kanamycin | pSG372 with 11x tetO array from Mahesri plasmid downstream of promoter, without 5' aptamer |
| pPC011 | pPC000 | Kanamycin | pPC000 with cytoplasmic Cypridina luciferase (CLuc) instead of FLuc, an SV40 signal at the start with a unique BamHI site, and a new unique XmaI site at the end of the tetR CDS. CLuc was found not to function in P. falciparum. High expression of SV40_TetR was found to be toxic in absence of inducer. |
| pPC012 | pPC011 | Kanamycin | pPC011 with 8x tetO array from Mahesri plasmid downstream of promoter. CLuc was found not to function in P. falciparum. High expression of SV40_TetR was found to be toxic in absence of inducer. |

| | | | |
|---|---|---|---|
| pPC013 | pPC011 | Kanamycin | pPC011 with 8x compact tetO array downstream of promoter, unnecessary XhoI cut site before operator array (besides after). CLuc was found not to function in P. falciparum. High expression of SV40_TetR was found to be toxic in absence of inducer. |
| pPC023 | pPC011 | Kanamycin | pPC011 with 8x compact tetO array downstream of promoter (same as pPC013 but without unnecessary XhoI cut site before operator array). CLuc was found not to function in P. falciparum. High expression of SV40_TetR was found to be toxic in absence of inducer. |
| pPC029 | pPC013 | Kanamycin | pPC013 with full-length CLuc (including the wild-type secretion signal). CLuc was found not to function in P. falciparum. High expression of SV40_TetR was found to be toxic in absence of inducer. |
| pPC030 | pPC011 | Kanamycin | pPC011 with FLuc (including ATG) instead of CLuc. High expression of SV40_TetR was found to be toxic in absence of inducer. |
| pPC031 | pPC012 | Kanamycin | pPC012 with FLuc (including ATG) instead of CLuc. High expression of SV40_TetR was found to be toxic in absence of inducer. |
| pPC032 | pPC023 | Kanamycin | pPC023 with FLuc (including ATG) instead of CLuc. High expression of SV40_TetR was found to be toxic in absence of inducer. |
| pPC038 | pPC032 | Kanamycin | pPC032 with lacI instead of tetR and 8x compact operator array with a mix of lacO and Osys sequences downstream of promoter; requires IPTG to clone, array is unstable—grow at 30 ºC to be safe. A more stable array was developed later (pPC089). |
| pPC0 | pPC032 | Kanamycin | pPC032 with cymR instead of tetR and 8x compact CymR operator array downstream of promoter |
| pPC040 | pPC032 | Kanamycin | pPC032 with phlF instead of tetR and 8x compact PhlF operator array downstream of promoter (contains ATGs and stop codons in operator array). Operators mutated to have no stop codons and a T2A peptide added between operator array and CLuc. |
| pPC041 | pPC032 | Kanamycin | pPC032 with phlF instead of tetR and 8x compact PhlF operator array downstream of promoter (contains ATGs and stop codons in operator array). Operators mutated to have no start codons. |
| pPC070 | pPC032 | Kanamycin | pPC032 with no SV40 |
| pPC071 | pPC032 | Kanamycin | pPC032 with mRPL2 promoter on SV40_TetR-RLuc-BSD instead of HSP86 promoter |
| pPC072 | pPC032 | Kanamycin | pPC032 with TetR-Sir2A instead of SV40_TetR |
| pPC082 | pPC031 | Kanamycin | pmRPL2-SV40_TetR_Sir2a operating on pCAM-extended 8x tetO |
| pPC083 | pPC082 | Kanamycin | pmRPL2-SV40_TetR_Sir2a operating on pCAM-extended 11x tetO |
| pPC088 | pPC038 | Kanamycin | pPC038 but with 4x extended lacO array (1 Osys) |
| pPC089 | pPC038 | Kanamycin | pPC038 but with 8x extended lacO array (1 Osys) |
| pPC094 | pPC071 | Kanamycin | pPC071 with PfCRT promoter instead of PfCAM |
| pPC095 | pPC071 | Kanamycin | pPC071 with PcDT (AMA1) promoter instead of PfCAM |
| pPC096 | pPC071 | Kanamycin | pPC071 with BiP1_PB (HSP70) promoter instead of PfCAM |
| pPC097 | pPC071 | Kanamycin | pPC071 with EF1alpha (PF3D7_1357000/PF3D7_1357100) promoter instead of PfCAM |
| pPC098 | pPC071 | Kanamycin | pPC071 with mNeonGreen instead of FLuc |
| pPC099 | pPC089 | Kanamycin | pPC089 with mNeonGreen instead of FLuc |

| | | | |
|---|---|---|---|
| pPC101 | pSN150 | Chloramphenicol | pPC089 with no FLuc moved into a linear pSN150 context (derived from pJAZZ, see Nasamu et al. 2020). Base plasmid for native gene regulation. |
| pPC111 | pPC089 | Kanamycin | pPC089 with PfCRT promoter instead of PfCAM |
| pPC112 | pPC089 | Kanamycin | pPC089 with PcDT (AMA1) promoter instead of PfCAM |
| pPC113 | pPC089 | Kanamycin | pPC089 with BiP1_PB (HSP70) promoter instead of PfCAM |
| pPC114 | pPC089 | Kanamycin | pPC089 with EF1alpha (PF3D7_1357000/PF3D7_1357100) promoter instead of PfCAM |
| pPC119 | pPC030 | Kanamycin | pPC030 with the SV40 NLS tag removed |
| pPC120 | pPC071 | Kanamycin | pPC071 with no tetO array |
| pPC121 | pPC089 | Kanamycin | pPC089 with no lacO array |
| pPC126 | pPC121 | Kanamycin | pPC121 with 3' 8x lacO |
| pPC127 | pPC101 | Chloramphenicol | pPC101 with LHR and RR+RHR+gRNA inserted for PI3K control (PF3D7_0515300) |
| pPC129 | pPC101 | Chloramphenicol | pPC101 with LHR and RR+RHR (232 bp)+gRNA inserted for PI4K control (PF3D7_0509800) |
| pPC150 | pPC101 | Chloramphenicol | pPC101 with LHR and RR+RHR+gRNA (no HA tags) inserted for Kelch13 (PF3D7_1343700) |
| pPC151 | pPC101 | Chloramphenicol | pPC101 with LHR and RR+RHR+gRNA (with HA tags) inserted for Kelch13 (PF3D7_1343700) |

**Table S2.** DNA oligos used in this study.

| Name | Type | Use | Sequence |
|---|---|---|---|
| PC_009 | DNA Oligo | Sanger sequencing | tctctgcctggattgcttaag |
| PC_010 | DNA Oligo | Sanger sequencing | gcttgctagagcgattacc |
| PC_011 | DNA Oligo | Sanger sequencing | ataaggcgcgcctctag |
| PC_012 | DNA Oligo | Sanger sequencing | tatccaatggcccctttcc |
| PC_013 | DNA Oligo | Sanger sequencing | tctctagcagggccgag |
| PC_014 | DNA Oligo | Sanger sequencing | atctagacatttttcctaggtttattcg |
| PC_015 | DNA Oligo | Sanger sequencing | tttcgaataaacctaggaaaaatgtctag |
| PC_016 | DNA Oligo | Sanger sequencing | gtatgccgccattattacgac |
| PC_017 | DNA Oligo | Sanger sequencing | gcatggtttgaacttcttaatttacc |
| PC_018 | DNA Oligo | Sanger sequencing | gaagatgcacctgatgaaatgg |
| PC_019 | DNA Oligo | Sanger sequencing | agggctaaaccggtccat |
| PC_020 | DNA Oligo | Sanger sequencing | gagctcgttttcgacactg |
| PC_021 | DNA Oligo | Sanger sequencing | attacgctcgtcatcaaaatcac |
| PC_022 | DNA Oligo | Sanger sequencing | cgtgagtcttttccttacccat |
| PC_023 | DNA Oligo | Sanger sequencing | accccgtagaaaagatcaaagg |
| PC_024 | DNA Oligo | Sanger sequencing | gatttttgtgatgctcgtcagg |
| PC_025 | DNA Oligo | Sanger sequencing | gcccgagctcgaattcc |
| PC_026 | DNA Oligo | Sanger sequencing | ggcatgctatttgatgaattaactac |
| PC_027 | DNA Oligo | Sanger sequencing | gtttacaactcggactttccg |
| PC_028 | DNA Oligo | Sanger sequencing | aaaatggaacaactttaccgacc |
| PC_029 | DNA Oligo | Sanger sequencing | cgggaggtagatgagatgtg |
| PC_030 | DNA Oligo | Sanger sequencing | gtgatagagacgacggaagg |
| PC_031 | DNA Oligo | Sanger sequencing | cagggcgtatctcttcatagc |
| PC_032 | DNA Oligo | Sanger sequencing | gggcgcgccttatatgtg |
| PC_033 | DNA Oligo | PCR, Gibson assembly | tctttatgtttttggcgtcttcatgttctcgagccatctctagcagcgggcagg |
| PC_034 | DNA Oligo | PCR, Gibson assembly | gtccgtatcgacctttctctgcctggattgcttaagctctagcagcgggcagg |
| PC_035 | DNA Oligo | PCR, Gibson assembly | atattttataataataaatacctaatagaaatatatcttaaggatagggaccttccgtcg |
| PC_036 | DNA Oligo | PCR, Gibson assembly | tataataataaatacctaatagaaatatatcttaagtcactgatagggaccacctgtctc |

| | | | |
|---|---|---|---|
| PC_037 | DNA Oligo | PCR, Gibson assembly | tatattttataataataaatacctaatagaaatatatcttaagccgtctcccctgctaga |
| PC_038 | DNA Oligo | PCR, Gibson assembly | ttttataataataaatacctaatagaaatatatcttaagcattgctagagttgtccccttc |
| PC_041 | DNA Oligo | Sanger sequencing | ccttccgtcgtctctatcac |
| PC_044 | DNA Oligo | PCR, Gibson assembly | gaaagtgggtctcccgggagtggtgaaggtagaggatcattattaac |
| PC_045 | DNA Oligo | PCR, Gibson assembly | ttataatatttttaatctattattaaataaatttaatggaccggtttagccctcccacac |
| PC_046 | DNA Oligo | PCR, Gibson assembly | ataaatacctaatagaaatatatcttaagctcgagatggactgtccttacgaacctgatc |
| PC_047 | DNA Oligo | PCR, Gibson assembly | tgtgagtacataaatatattatataaactagtTCAtttgcattcatctggtacttctagg |
| PC_048 | DNA Oligo | Anneal-extension, Gibson assembly | cactttacttttatctaatctggatccgaccttccgcttcttctttggggccatttttcc |
| PC_049 | DNA Oligo | Anneal-extension, Gibson assembly | tcttcccacatttcgaataaacctaggaaaaatggccccaaagaagaagcggaaggtcgg |
| PC_051 | DNA Oligo | Sanger sequencing | gacatactatcagatggactgtgt |
| PC_052 | DNA Oligo | Sanger sequencing | tgaccgtgtcgaacgatg |
| PC_053 | DNA Oligo | Sanger sequencing | gagaagcacctgtcggc |
| PC_054 | DNA Oligo | Sanger sequencing | aatgtaggctgctctacacc |
| PC_055 | DNA Oligo | Sanger sequencing | cctaggtttattcgaaatgtggg |
| PC_056 | DNA Oligo | Sanger sequencing | catccgtttcctttgttctgg |
| PC_057 | DNA Oligo | Sanger sequencing | acacagtccatctgatagtatgtc |
| PC_058 | DNA Oligo | Sanger sequencing | cccacatttcgaataaacctagg |
| PC_059 | DNA Oligo | PCR, Gibson assembly | agaagaagcggaaggtcggatccagattagataaaagtaaagtgattaacagcgcattag |
| PC_060 | DNA Oligo | PCR, Gibson assembly | atgatcctctaccttcaccactcccgggag |
| PC_061 | IDT Ultramer | Anneal-extension, Gibson assembly | tatattttataataataaatacctaatagaaatatatcttaagctcgagtctctatcactgatagggaAGTtctctatcactgatagggaCTtctctatcactgatagggaTACtctctatcactgatagggaGTTtctctatcactgatagggaGCtctctatcactgatagggaCGtctctatcactgatagggaGCT |
| PC_062 | IDT Ultramer | Anneal-extension, Gibson assembly | gttcgtaaggacagtccatctcgagtccctatcagtgatagagaAGCtccctatcagtgatagagaCGtccctatcagtgatagagaGCtccctatcagtgatagagaAACtccctatcagtgatagagaGTAtccctatcagtgatagagaAGtccctatcagtgatagagaACTtccctatcagtgatagagactcga |
| PC_063 | DNA Oligo | PCR, Gibson assembly | gatcaggttcgtaaggacagtccatctcgagccatctctagcagcg |
| PC_064 | IDT Ultramer | PCR, Gibson assembly | caatggcccctttccgggcgcgcctagtagttgagtgattctatatacatatacataaataaattac |
| PC_066 | IDT Ultramer | PCR, Gibson assembly | caatggcccctttccgggcgcgccATTAATGAGGTGTGTTGGGAAACAGAAGAAAAAAAC |
| PC_068 | DNA Oligo | PCR, Gibson assembly | caatggcccctttccgggcgcgccTTTTTTAATGTATGACATGTGTTGTAAATTTTGAGC |

| | | | |
|---|---|---|---|
| PC_070 | DNA Oligo | PCR, Gibson assembly | caatggcccctttccgggcgcgccTTTAACGGTTCACCCCTCTTAACC |
| PC_072 | IDT Ultramer | Anneal-extension, Gibson assembly | aagatttatattttataataataaatacctaatagaaatatatcttaagtctctatcactgatagggaAGTtctctatcactgatagggaCTtctctatcactgatagggaTACtctctatcactgatagggaGTTtctctatcactgatagggaGCtctctatcactgatagggaCGtctctatcactgatagggaGCT |
| PC_073 | IDT Ultramer | Anneal-extension, Gibson assembly | gttcgtaaggacagtccatctcgagtccctatcagtgatagagaAGCtccctatcagtgatagagaCGtccctatcagtgatagagaGCtccctatcagtgatagagaAACtccctatcagtgatagagaGTAtccctatcagtgatagagaAGtccctatcagtgatagagaACTtccctatcagtgatagagacttaa |
| PC_080 | IDT Ultramer | Anneal-extension, Gibson assembly | gatttatattttataataataaatacctaatagaaatatatcttaagaacaaacagacaatctggtctgtttgtaAGTaacaaacagacaatctggtctgtttgtaCTaacaaacagacaatctggtctgtttgtaTACaacaaacagacaatctggtctgtttgtaGTTaacaaacagacaatctggtctgtttgtaGC |
| PC_081 | IDT Ultramer | Anneal-extension, Gibson assembly | gttcgtaaggacagtccatctcgagtacaaacagaccagattgtctgtttgttAGCtacaaacagaccagattgtctgtttgttCGtacaaacagaccagattgtctgtttgttGCtacaaacagaccagattgtctgtttgttAACtacaaacagaccagattgtctgtttgttGTAtacaaacagaccagattgtctg |
| PC_084 | IDT gBlock gene fragment | Gibson assembly | atcctctaccttcaccactcccgggGCTGGCTCTCTGGGTGCCGGGGCACACGCCGTTGATCAGCAGAAAGGTGAACTCCTCAATATCCTGTTCCACGGTCAGCTGCTCGGTCAGCAGTCTGTACCAGCAGAAGCCGAAGATCATATCCAGCAGCAGTTCCCGGTTGGTGTCCTTGGGCAGCTCGCCGTTGCTGATGGCGTTTTCCACCAGTTTCTTGGGCATCTCTCTCCGCCGTTCCATGAACTGGTCCTTCAGCTGTGTCAGGGTGGCAGGGTCCAGCTGGGCTTCAGCGATCACGCATCTGAAGGCTTCGCCGCAGATGGTTTCCCGCCACACTTTCCACAGGTTCCGCAGCAGGAAGTCCAGATCGGCCTTGAAGCTGCCCAGGTCGGGGAACTTTCGGACCTGCTCGCTCTCGTTCTCGTACACCTCTGCGATCAGGGCGGCCTTGTTTGTCCACCATCGGTAGATGGTGGGCTTGGACGCGCCAGCCCTTCTAGCCACAGACTCGATGCTCAGGCCGCTGTAGCCGCACTCTTTCAGGATCTCGATGGTGCTGGTCAGGATGGCCTTGTGGGTGTGGGGGCTTCTCAGGCTGCCGATAGAGCTCCTAGAGGGGGTCCGGGCggatccgaccttccgcttcttcttt |
| PC_086 | IDT gBlock gene fragment | Gibson assembly | atcctctaccttcaccactcccgggctgcccgctttccagtcgggaaacctgtcgtgccagctgcattaatgaatcggccaacgcgcggggagaggcggtttgcgtattgggagccagggtggtttttcttttcaccagtgagacgggcaacagctgattgcccttcaccgcctggccctgagagagttgcagcaagcggtccacgctggtttgccccagcaggcgaaaatcctgtttgatggtggtcaacggcgggatataacatgagctgtcttcggtatcgtcgtatcccactaccgaaatatccgcaccaacgcgcagcccggactcggtaatggcacgcatagcgcccagcgccatctgatcgttggcaaccagcatcgcagtgggaacgatgccctcattcagcatttgcatggtttgttgaaaaccggacatggcactccagtcaccttcccgttccgctatcggctgaatttgattgcgagtgagatatttatgccagccagccagacgcagacgcgctgagacagaacttaatggacccgctaacagcgcgatttgctggtgtcccaatgcgaccagatgctccacgcccagtcgcgtaccgtcttcatgggagaaaataatactgttgatgggtgtctggtcagagacatcaagaaataacgccggaacattagtgcaggcagcttccacagcaatagcatcctggtcatccagcggatagttaatgatcagcccactgacacgttgcgcgagaagattgtgaaccgccgctttacaggcttcgacgccgcttcgttctaccatcgacaccaccacgctggcacccagttgatcggcgcgagatttaatcgccgcgacaatttgcgacggcgcgtgcagggccagactggaggtggcaacgccaatcagcaacgactgtttgcccgccagttgttgtgccacgcggttgggaatgtaattcagctccgccatcgccgcttccactttttcccgcgttttcgcagaaacgtggctggcctggttcaccacgcgggaaacggtctgataagagacaccggcatactcagcgacatcgtataacgtaactggtttggatccgaccttccgcttcttctttg |
| PC_087 | IDT gBlock gene fragment | Gibson assembly | atcctctaccttcaccactcccggggcgcttgaatttcgcgtaccgctctcgcgcaatttcgagtgtcgagttcctgacacgctcaaagcgttctttgtccttctgccataggctacgaacagcaagcccacgcaccgaattgaatatcaaccaaaggatgtcttctgcatcatcacgcgaaagaccacggctcaccagaacaccaagccacatatcctcgacgacaaaccgattcctctctaccgtgcgctgaataccctcgcgtaacgctgggtcccggtcggcagccacaatcaaatcaaggctgatagagaagtcatcgtcgaggaaaaaattcggcggcgtcgtccagcatttgctggatgacatcatcctctggcttcaatttcgctaatcgagcccgactgcgttcggtgatctgttcgtaaagccattcaaaagtggcaagcagaagct |

| | | | |
|---|---|---|---|
| | | | caagctttgtcgggaaatgatggctctgcgctcctctggagacaccagcagcaccgggcacatctgcgatacggaatcccgcgtaaccttttcccgtaaaccccccagagccgctgcaatcaacttgccttgggtctccatagcccgctctgcctgtgttcttctctttggactggatccgaccttccgcttcttcttt |
| PC_089 | DNA Oligo | Sanger sequencing | tcctgcggcctctactc |
| PC_090 | DNA Oligo | PCR, Gibson assembly | aagatttatattttataataataaatacctaatagaaatatatcttaag |
| PC_094 | DNA Oligo | Anneal-extension, Gibson assembly | cactttacttttatctaatctggatcccatcctaggtttattcgaaatgtgggaaga |
| PC_095 | DNA Oligo | Anneal-extension, Gibson assembly | tcttcccacatttcgaataaacctaggatgggatccagattagataaaagtaaagtg |
| PC_096 | IDT Ultramer | PCR, Gibson assembly | cctaatagaaatatatcttaagtctgttatccgctcacatctcataattgtgagcgctcacaattttCTttgttatccgctcacaaTACaattgtgagcgctcacaattACGtgtgagcggataacattgcaaattgtgagcgctcacaattGTTttgtgagcggataacaagaGCTaattgtgagcgctcacaattctc |
| PC_097 | IDT Ultramer | PCR, Gibson assembly | aggacagtccatctcgagaattgtgagcgctcacaattAGCtcttgttatccgctcacaaAACaattgtgagcgctcacaatttgcaatgttatccgctcacaCGTaattgtgagcgctcacaattGTAttgtgagcggataacaaAGaaaattgtgagcgctcacaattatgagatgtgagcggataacagacttaaga |
| PC_100 | IDT Ultramer | Anneal-extension, Gibson assembly | agatttatattttataataataaatacctaatagaaatatatcttaagATGATACGAAACGTACCGTATCGTTACGGTAGTATGATACGAAACGTACCGTATCGTTACGGTCATATGATACGAAACGTACCGTATCGTTACGGTTACATGATACGAAACGTACCGTATCGTTACGGTGTTATGATACGAAACGTACCGTA |
| PC_101 | IDT Ultramer | Anneal-extension, Gibson assembly | gatcctctaccttcaccactcccACCGTAACGATACGGTACGTTTCGTATCATAGCACCGTAACGATACGGTACGTTTCGTATCATACGACCGTAACGATACGGTACGTTTCGTATCATGTGACCGTAACGATACGGTACGTTTCGTATCATAACACCGTAACGATACGGTACGTTTCGTATCATGTAACCGTAACGATA |
| PC_104 | IDT Ultramer | Anneal-extension, Gibson assembly | aagatttatattttataataataaatacctaatagaaatatatcttaagACGATACGAAACGTACCGTATCGTTAAGGTAGTACGATACGAAACGTACCGTATCGTTAAGGTCTACGATACGAAACGTACCGTATCGTTAAGGTTACACGATACGAAACGTACCGTATCGTTAAGGTGTTACGATACGAAACGTACCGTA |
| PC_105 | IDT Ultramer | Anneal-extension, Gibson assembly | gttcgtaaggacagtccatctcgagACCTTAACGATACGGTACGTTTCGTATCGTAGCACCTTAACGATACGGTACGTTTCGTATCGTGGACCTTAACGATACGGTACGTTTCGTATCGTACACCTTAACGATACGGTACGTTTCGTATCGTAACACCTTAACGATACGGTACGTTTCGTATCGTGTAACCTTAACGATA |
| PC_111 | DNA Oligo | PCR, Gibson assembly | cgttgcattagtctactgcgccactgttcattgtcaggactgtccttacgaacctgatcc |
| PC_113 | DNA Oligo | PCR, Gibson assembly | gtagaccccattgtgagtacataaatatattatataaactagt |
| PC_114 | DNA Oligo | PCR, Gibson assembly | GATAGGGAGCTTCTCTATCACTGATAGGGACTCGAGATGGAAGACGCCAAAAACATA |
| PC_117 | DNA Oligo | PCR, Gibson assembly | TTTTGTACAATTTATAACAAGTACATTAATTTGAAAAATTATC |
| PC_119 | IDT Ultramer | PCR, Gibson assembly | ccctatcagtgatagagacttaagGTTTAGTATATTAATATATATGTATATATATATATATATATATTTTTTTTTTTTCTATTTATATTTATTTATATAAATCAG |
| PC_122 | DNA Oligo | PCR, Gibson assembly | gaGCTaattgtgagcgctcacaattctcgagatggaagacgccaaaaac |
| PC_123 | DNA Oligo | PCR, Gibson assembly | aacaaacagacaatctggtctgtttgtactcgagatggaagacgccaaaaac |

| | | | |
|---|---|---|---|
| PC_126 | IDT Ultramer | PCR, Gibson assembly | tatattttataataataaatacctaatagaaatatatcttaagctcgagatggaagacgccaaaaacataaagaaagg |
| PC_127 | DNA Oligo | PCR, Gibson assembly | tcagtgatagagaACTtccctatcagtgatagagacttaag |
| PC_129 | DNA Oligo | PCR, Gibson assembly | AAATTAATGTACTTGTTATAAATTGTACAAAActtaagtctctatcactgatagggaAGT |
| PC_130 | DNA Oligo | PCR, Gibson assembly | ATTTTTTTTATTATCATATATATAATTCAAAActtaagtctctatcactgataggg aAGT |
| PC_131 | IDT Ultramer | PCR, Gibson assembly | cctatcagtgatagagaACTtccctatcagtgatagagacttaagTTTTGAATTATATATATGATAATAAAAAAAATTAAATAAATAAATATAAATATGTTTTATTTATATTTTAAAATATATATATATATAAGG |
| PC_178 | DNA Oligo | PCR, Gibson assembly | ggtggaggttctggtggaggatctCCTGCAGGTATGGGTAATTTAATGATTTCCTTTTTG |
| PC_179 | DNA Oligo | PCR, Gibson assembly | caagaagggcggaaagtccgagttgtaaactagtCAGAATGTGTCCACATTAGGAACATG |
| PC_187 | IDT Ultramer | PCR, Gibson assembly | ctggatccgaccttccgcttcttctttggggccatttttcctaggTATATTGTATAATATTAACAAAAAAAAATAATAATAATAA |
| PC_199 | IDT Ultramer | PCR, Gibson assembly | caccacatgttaataatgatcctctaccttcaccactcccgggCATTATTTTCTTATTTTTTTCACTTGACC |
| PC_200 | IDT Ultramer | PCR, Gibson assembly | ctttttattatattttttttcttcccacatttcgaataaacctaggatgggatccagattagataaaagtaaagt |
| PC_207 | IDT Ultramer | PCR, Gibson assembly | tatataaattgataatatataaaattaatcacatataaggcgcgccACTTACATTATTTATTTATATAAAAAGTTAATTATAATAAAATATAC |
| PC_208 | DNA Oligo | Sanger sequencing | aaattaatcacatataaggcgcg |
| PC_215 | DNA Oligo | Sanger sequencing | ttcgatccgcttcattctctc |
| PC_217 | IDT Ultramer | Anneal-extension, Gibson assembly | aagatttatattttataataataaatacctaatagaaatatatcttaagATTTATATTATCTCGATTAAAAAATTGGTATCCGCTCACAATTCCAATTTTTTTTAAAATTGTTATCCGCTCACAATACCACAGAACGTAAAATTGGTATACGCTCACAATACCACAATATAAAaattgtgagcgctcacaattTCGACAc |
| PC_218 | IDT Ultramer | Anneal-extension, Gibson assembly | tggcgccgggcctttctttatgtttttggcgtcttccatctcgagctctagcagcgggcagggTGTCGAaattgtgagcgctcacaattTTTATATTGTGGTATTGTGAGCGTATACCAATTTTACGTTCTGTGGTATTGTGAGCGGATAACAATTTTAAAAAAAATTGGAATTGTGAGCGGATACCAATTTTTTAATCG |
| PC_219 | IDT Ultramer | Anneal-extension, Gibson assembly | aagatttatattttataataataaatacctaatagaaatatatcttaagccgtctcccctgctAATTGTTATCCGCTCACAATTCCACATTAGTCATAATTGGTATTCGCTCACAATACCACAATATTACTTAAATTGTTATACGCTCACAATTCCTCACATATCATAAATTGGTATCCGCTCACAATACCAATttaagA |
| PC_220 | IDT Ultramer | Anneal-extension, Gibson assembly | GCGGATACCAATTTTTTAATCGAGATAATATAAATcttaaATTGGTATTGTGAGCGGATACCAATTTATGATATGTGAGGAATTGTGAGCGTATAACAATTTAAGTAATATTGTGGTATTGTGAGCGAATACCAATTATGACTAATGTGGAATTGTGAGCGGATAACAATTagcaggggagacggcttaagatatatttc |
| PC_226 | DNA Oligo | PCR, Gibson assembly | tgtatatagaatcactcaactactaggcgcgccACTTACATTATTTATTTATATAAAAAG |
| PC_227 | DNA Oligo | PCR, Gibson assembly | CTGTTTCCCAACACACCTCATTAATggcgcgccACTTACATTATTTATTTATATAAAAAG |
| PC_229 | DNA Oligo | PCR, Gibson assembly | GGGTTAAGAGGGGTGAACCGTTAAAggcgcgccACTTACATTATTTATTTATATAAAAAG |
| PC_230 | DNA Oligo | Sanger sequencing | gcgttttcgcagaaacgtgg |

| | | | |
|---|---|---|---|
| PC_231 | IDT gBlock gene fragment | Gibson assembly | TAGGGAGCTTCTCTATCACTGATAGGGACTCGAGAATGGTTAGCAAGGGCGAAGAGGATAACATGGCCTCTCTCCCAGCGACACATGAGCTTCACATCTTCGGCTCTATCAACGGCGTCGATTTTGACATGGTAGGGCAGGGCACGGGCAATCCGAATGACGGATATGAAGAACTGAACCTGAAGTCCACAAAGGGTGACTTACAATTCTCCCCGTGGATTCTGGTCCCTCATATTGGGTATGGCTTCCATCAGTACCTCCCCTACCCTGACGGGATGAGCCCTTTCCAGGCCGCAATGGTCGATGGATCTGGGTATCAAGTCCACCGCACAATGCAGTTTGAAGACGGTGCGAGTCTTACTGTTAATTACCGCTACACCTACGAAGGAAGCCACATAAAAGGAGAGGCCCAGGTGAAGGGGACGGGATTCCCGGCTGACGGTCCCGTGATGACAAACTCGCTGACGGCTGCGGACTGGTGCAGATCGAAGAAGACTTACCCCAACGACAAAACGATCATCTCAACCTTCAAGTGGTCGTACACCACTGGAAATGGCAAACGATACCGGAGCACTGCGCGTACCACTTACACGTTTGCAAAACCAATGGCGGCTAACTACTTGAAAAACCAGCCGATGTACGTGTTCAGGAAGACGGAGCTCAAGCACTCTAAAACAGAGCTCAACTTCAAAGAGTGGCAAAAGGCATTTACGGATGTGATGGGCATGGACGAGTTGTATAAGTGAGGGCCCACTAGTTTATATAATATATTTATGTACTCACAATGGGGTCTACAA |
| PC_232 | DNA Oligo | PCR, Gibson assembly | ctcacaattTCGACAccctgcccgctgctagagctcgagaATGGTTAGCAAGGGCGAAGA |
| PC_235 | IDT Ultramer | PCR, Gibson assembly | cggtttaccgagctcttattggttttcaaacttcattgactgtgccggccggccgacaataaaaagattcttgttttcaagaacttg |
| PC_241 | DNA Oligo | Sanger sequencing | TGTCAAAATCGACGCCGTTG |
| PC_242 | DNA Oligo | Sanger sequencing | agcgcatgggtaaggaaggg |
| PC_243 | DNA Oligo | Sanger sequencing | cggtggttgaccagacaaacc |
| PC_256 | IDT Ultramer | PCR, Gibson assembly | ctggccgtcgttttacaacgtcgtgactgggaaaaccctggcgtcgagcgatcgcctctagcagcgggcagggTGTCG |
| PC_260 | DNA Oligo | PCR, Gibson assembly | ggcgcgccTTTTTTAATGTATGACATGTGTTGTAAATTTTGAGC |
| PC_264 | IDT Ultramer | PCR, Gibson assembly | GAGCGGATAACAATTagcaggggagacggcttaagtgttatatgtaagaaattaaaaaaaacaaaataataaatg |
| PC_265 | IDT Ultramer | PCR, Gibson assembly | TGAGCGGATAACAATTagcaggggagacggcttaagTTTTGTACAATTTATAACAAGTACATTAATTTG |
| PC_266 | IDT Ultramer | PCR, Gibson assembly | GAGCGGATAACAATTagcaggggagacggcttaagTTTTGAATTATATATATGATAATAAAAAAAATTAAATAAATAAATATAAATATG |
| PC_267 | IDT Ultramer | PCR, Gibson assembly | GAGCGGATAACAATTagcaggggagacggcttaagGTTTAGTATATTAATATATATGTATATATATATATATATATATATTTTTTTTTTTTC |
| PC_268 | DNA Oligo | PCR, Gibson assembly | tggaatactaaatatatatccaatggcccctttcc |
| PC_269 | DNA Oligo | PCR, Gibson assembly | aaaaatatataaattgataatatataaaattaatcacatataaggcgcgccGCATGAACG |
| PC_270 | IDT Ultramer | PCR, Gibson assembly | GAATAAACTAGTGCTCAAAATTTACAACACATGTCATACATTAAAAAAggcgcgccACTTACATTATTTATTTATATAAAAAG |
| PC_270 | IDT Ultramer | PCR, Gibson assembly | GAATAAACTAGTGCTCAAAATTTACAACACATGTCATACATTAAAAAAggcgcgccACTTACATTATTTATTTATATAAAAAG |
| PC_288 | DNA Oligo | PCR, Gibson assembly | aaggccaagaagggcggaaagtccgagttgtaaactagtccgtctcccctgctAATTGTT |
| PC_289 | DNA Oligo | PCR, Gibson assembly | gtagaccccattgtgagtacataaatatattatataagggcccctctagcagcgggcagg |
| PC_294 | DNA Oligo | PCR, Gibson assembly | tcaaacttcattgactgtgccggccggccTGCCCCTTCAAAGAAAAAATACATATTTATG |

| | | | |
|---|---|---|---|
| PC_295 | DNA Oligo | PCR, Gibson assembly | aacaagaatctttttattgtcggccggccTCTTCTTTGTGGTATATACTCTCCTATATAC |
| PC_296 | IDT gBlock gene fragment | Gibson assembly | gcgctcacaattTCGACAccctgcccgctgctagaggcgATGGGCAGCTACCCCTACGACGTACCAGACTACGCCGGAAGCTACCCATACGACGTTCCGGACTATGCAGGAAGCATGAAAATCAGATACGACAAGTGTAGCAGCACCAAAGATCTAAACTATTTCTTTCACCTAAAGCTGGGATTCTTTGTATGCTATAAGAACCACAACGACAAATATAGCTTCAAGAACAAGATTTTGCAAAAAAACGACACGATTCTCTTCTTCAAGAAGAAGAAGAAGTTCATGTACCTTCGCAAGAAGAAGAAAAAGAAAAAAAAGAAAATACTCATTCAGATTATACAAGAATATAATAAATATAATGAATATTTTAAATATAATTCAAATTTAGAAGGTAATCAAGGATTCAATAAAAAACCAGAAAAAAATAAAAACACAAAAGGAAATGTATATACTGATCATACAAATCAAAATGCAAAATCAAAAATATATAATTATGATATGAATGATGATTCTTATTCTAATTATGTGAACAACAATAATGTTTTTAGAATATCATCTTTTTTGATATTAAATAATGAATTTTTTGGATATCCCCTTCAATTTGTTTGTGAAACAGAAGGGAGAAGTAGGAACCATGAACATTATCCGGACGTACATGGGGATAATATTAAGTATAACAAATGTGATGATAATAAGTATcgccgtacgtaatacgactcactataggGTAGTTTAGGTCTTTTGTACgttttagagctagaaatagcaagttaaaataaggctagtccgttatcaacttgaaaaagtggcaccgagtcggtgctactctaaccccatcggccgtcttaggggtttttgtgcgatcgctcgacgccagggttttcccagtcacgacgt |
| PC_297 | DNA Oligo | PCR, Gibson assembly | ttcattgactgtgccggccggccTGATATGAGTTATCCAAAATTGAAACATATATTTTTC |
| PC_298 | DNA Oligo | PCR, Gibson assembly | tgaaaacaagaatctttttattgtcggccggccTTTTACCACAGGTTAAAAAAAAAGAAG |
| PC_303 | Twist gene fragment | Gibson assembly | attTCGACAccctgcccgctgctagaggcgATGGGCAGCTATCCGTATGACGTTCCGGACTATGCTGGATCCTATCCATACGACGTACCGGACTATGCGGGCAGCATGGAGGACGACAAGGACAACAACATTACGAAGGAGAACTTAAACGAGGTTGACATTGAGAACAGCCAGCACCCGATTATTGAGGAGAAGAACTTAAAGGAGGAGGGCGTTAAGGACGAGGGCGGCAAGGTTTTTTTAAAAGGTGAACCGATACATAAACATTTATACAAACATAGTTATAATACGCAAGTATATAAAGATGAAGTTACACATATGCGTTTAAGAAATAGAGAAAATAAAAATTTGAATGAAAATAATTCGGATGATAATAAAAAGGTAAATAAGAAGAATGGGTGTGTTGATATTAATATGAATGAACATACTAATAAAAAGGAAGATGGTGATGATGATGATGGTGATGcgccgtacgtaatacgactcactataggGAAGGAGTAAAAGATGAGGGgttttagagctagaaatagcaagttaaaataaggctagtccgttatcaacttgaaaaagtggcaccgagtcggtgctactctaaccccatcggccgtcttaggggtttttgtgcgatcgctcgacgccagggttttcccagtcacgacgt |
| PC_338 | Twist gene fragment | Gibson assembly | cgctcacaattTCGACAccctgcccgctgctagaggcgATGGGCAGCTATCCGTATGACGTTCCGGACTATGCGGGATCGTATCCATACGATGTACCCGACTACGCGGGCAGCGGCAGCGGCATGGAGGGCGAGAAGGTTAAGACGAAGGCGAACAGCATTAGCAACTTTAGCATGACGTATGACCGCGAGAGCGGTGGTAACAGCAATAGTGATGATAAAAGCGGAAGTAGTAGCGAGAATGATTCTAATTCATTTATGAATCTAACTAGTGATAAAAATGAGAAAACGGAAAATAATAGTTTCCTTTTAAATAATAGTAGTTATGGAAATGTTAAAGATAGCCTATTAGAATCCATTGATATGAGTGTATTAGATTCGAACTTTGATAGTAAAAAAGATTTTTTACCAAGTAATTTATCAAGAACATTTAATAATATGTCTAAAGATAATATAGGcgccgtacgtaatacgactcactataggACGTATGATAGGGAATCTGGgttttagagctagaaatagcaagttaaaataaggctagtccgttatcaacttgaaaaagtggcaccgagtcggtgctactctaaccccatcggccgtcttaggggtttttgtgcgatcgctcgacgccagggttttcccagtcacgacgttgtaaaacg |
| PC_350 | Twist gene fragment | Gibson assembly | gagcgctcacaattTCGACAccctgcccgctgctagaggcgATGGGCAGCTATCCGTATGACGTTCCGGACTATGCGGGGTCATACCCCTACGATGTCCCCGACTATGCGGGCAGCGGCAGCGGCATGGAGGACAACAACATGAACAACGAGCAGAAGTTTAACCCGAACATGTTAGGCATGGAGAACAACGTTAGCCAGGAGCCGGCGGCGGAGACGGTTGTTGTTGAGCCGAAGAAGAAGGTTGGCCGCCCGCGCGGCGCGAGCAGCAAC |

| | | | |
|---|---|---|---|
| | | | AACAAGGTTATTAAGAAGGAGGAGAAGGTTAGCACGAGCAGCAG<br>CGGCTATCCGGGCGTTAGCTGGAACAAGCGTATGTGTGCGTGGC<br>TAGCTTTTTTCTATGATGGTGCATCAAGAAGAAGTAGAACATTTCA<br>TCCAAAACACTTTAATATGGATAAAGAAAAAGCAAGACTAGCAGC<br>AGTAGAATTTATGAAGACTGTTGAGAATAATGGTAGAAAAAAATCT<br>GGAAAAGGTAAAGGTGGTAGAAGTAAAAGCAAACAATTAAATGAT<br>GAACATTTCAATGCATTACACAATAGTGATCTTGGAAACAGTATGA<br>ATGGTATGAATGCTCATAATAATTTACACATGCAGTTGATGTCTAT<br>GAATCCAGCTTTTTATATGCAGAGTTTAAACAGAAATTTTAATATG<br>TCAGCAAACGAAAGAAATAATATGTTTAACTCTTTGAATAATCATT<br>CTGGATTAAGTGGACCACTTcgccgtacgtaatacgactcactataggCGCTT<br>ATTCCAACTAACACCgttttagagctagaaatagcaagttaaaataaggctagtccgtta<br>tcaacttgaaaaagtggcaccgagtcggtgctactctaaccccatcggccgtcttaggggtttttgt<br>gcgatcgctcgacgccagggttttcccagtcacgacgttgtaaaacgacggccagtgaattg |
| PC_355 | DNA Oligo | RT-qPCR for PI3K | tggggtgcacaaatccagtcgt |
| PC_356 | DNA Oligo | RT-qPCR for PI3K | tgctgcccggataattcaccct |
| PC_357 | DNA Oligo | RT-qPCR for PI4K | ggcgataaactgatgccgtcgc |
| PC_358 | DNA Oligo | RT-qPCR for PI4K | tgccacaggtttatcatgcgacg |
| PC_375 | DNA Oligo | RT-qPCR for actin (PFL2215w, Salanti et al. 2003) | AGCAGCAGGAATCCACACA |
| PC_376 | DNA Oligo | RT-qPCR for actin (PFL2215w, Salanti et al. 2003) | TGATGGTGCAAGGGTTGTAA |
| PC_377 | IDT Ultramer | PCR, Gibson assembly | accgagctcttattggttttcaaacttcattgactgtgccggccggccAAAAATTTTATAAA<br>TTAAATTTAGAAGAAAAATTTTGTTATATATAATAATAATATTAATAA<br>ATTTTCTGATCA |
| PC_378 | IDT Ultramer | PCR, Gibson assembly | gacaagttcttgaaaacaagaatctttttattgtcggccggccctttttctccttccatAATTATAA<br>TTTAATTAACAAAAACATAATAAATGAATG |
| PC_381 | DNA Oligo | RT-qPCR for FLuc | tatcaggtggcccccgctgaat |
| PC_382 | DNA Oligo | RT-qPCR for FLuc | ctccaaaacaacaacggcggcg |
